# A neural circuit basis for reward-induced suppression of fear generalization

**DOI:** 10.1101/2024.09.05.611495

**Authors:** Mi-Seon Kong, Yong S. Jo, Ekayana Sethi, Gyeong Hee Pyeon, Larry S. Zweifel

## Abstract

How positive and negative affective stimuli interact in the brain to influence behavioral outcomes remains poorly understood. Here, we show that recall of a positive reward-associated conditioned stimulus (CS_Rew_+) can prevent or reverse fear generalization in mice. Modification of generalized fear by recall of a CS_Rew_+ is dependent on the midbrain dopamine system and the regulation of discriminatory threat encoding by the central amygdala (CeA). Precisely timed, transient elevations in dopamine and activation of dopamine D2 receptors in the CeA are necessary to reverse threat generalization and non-discriminatory threat encoding in the CeA. Recall of a positive association is also effective at enhancing the extinction of a conditioned threat response in a dopamine dependent manner. These data demonstrate that recall of a positive experience can be an effective means to suppress generalized fear and show that dopamine projections to the CeA are an important neural substrate for this phenomenon.

## Introduction

The ability to learn stimulus-outcome associations is predicated on effective stimulus discrimination. Stress caused by uncertainty, and vice versa, can impair discriminatory fear learning resulting in maladaptive generalization^1–3^. Negative affective stimuli can potently suppress positive reinforcement^4^, but the extent to which positive affective stimuli can impact discriminatory fear learning is less clear.

In generalized fear-related disorders, like post-traumatic stress disorder, the most efficacious therapy is exposure-based treatment where recall of the most salient features of trauma-related memory is repeatedly evoked to neutralize or extinguish those memories^5^. In human subjects, positive emotions have been shown to have an inhibitory effect on fear generalization^6^, and positive affect has been shown to enhance fear extinction^7^. In rodents, counterconditioning in which presentation of a fear-associated conditioned stimulus (CS_Fear_+) is subsequently paired with a reward unconditioned stimulus (US_Rew_) results in persistent reductions in the fear memory that is associated with immediate early gene induction in reward circuitry^8^. These findings suggest that stimuli associated with positive affect can have a potent influence on fear-related memories, but the exact nature of how a positive stimulus affects the encoding of fear to exact these outcomes is not clear.

The ventral tegmental area (VTA) dopamine system and the CeA have emerged as two key nodes for regulating behavioral responses to both positive and negative affective stimuli^9–12^. There are numerous connections between the CeA and midbrain dopamine systems^13–16^. Among these, the VTA dopamine neurons are the most broadly implicated in associative learning^17^ and are potent modulators of discriminatory fear learning^12,13,18^, suggesting that the connections between the VTA and CeA may be central to the interactions between positively and negatively valenced stimuli^19^. Here, we sought to determine whether positive reinforcement learning could be leveraged as a means to suppress or reverse fear generalization, or enhance extinction of a fear memory in a manner that is dependent on dopaminergic modulation of threat encoding in the CeA. We find that precisely timed delivery of a CS_Rew_+ at the onset of a fear CS can prevent and reverse fear generalization. This effect is associated with a CS_Rew_+ evoked change in dopamine release in the CeA during fear reconditioning and a shift from non-discriminatory to discriminatory fear encoding by neurons of the CeA. Mechanistically, dopamine release from the VTA and dopamine d2 receptor expression in the CeA are required for CS_Rew_+ modification of generalized fear memories. Consistent with this observation, the CS_Rew_+ modifies discriminatory fear encoding by Drd2-expressing neurons in the CeA. Finally, the timed delivery of the CS_Rew_+ in place of an expected fear US during extinction can enhance the extinction of the fear memory in a dopamine D2 receptor-dependent manner. Collectively, these data provide evidence for a neural substrate through which a reward associated stimulus can potently modify generalized fear states.

### A reward conditioned stimulus can prevent and reverse generalized fear

We first sought to establish whether a CS associated with positive reinforcement can impact threat generalization and whether this is selective for the reward-predictive cue. To achieve this, we utilized a Pavlovian reward reinforcement task (**Extended Data Fig. 1a**) in which a 10 s tone (CS_Rew_+) co-terminated with the delivery of a food reinforcer (US_Rew_; 25-CS/US pairings). Interleaved with the CS/US pairings, a second tone was played (10 s) an equal number of times but was not paired with reward delivery (CS_Rew_-). To assess the efficacy of conditioning, we monitored discriminatory conditioned responses (head entries during CS_Rew_+ presentation) and recorded dopamine neuron activity during CS_Rew_+ or CS_Rew_- presentations. Dopamine neurons were isolated during recording by injecting mice expressing Cre recombinase under the control of the dopamine transporter locus (DAT-Cre)^20^ with a virus for conditional expression of the stimulatory light-activated channel rhodopsin^21^ (AAV-FLEX-ChR2-mCherry) and implanting an optical fiber and electrodes over the VTA (**Extended Data Fig. 1b**). During the positive reinforcement task, mice learned to discriminate between the CS_Rew_+ and CS_Rew_- (**Extended Data Fig. 1c**). Photosensitive dopamine neurons (**Extended Data Fig. 1d-g**) exhibited increased conditioned responses to the CS_Rew_+ and diminished responses to the CS_Rew_- (**Fig. 1a,b**; CS response) which coincided with reduced responding to the reward US (**Fig. 1a,b**; Reward response), as predicted^22^.

**Figure 1.**
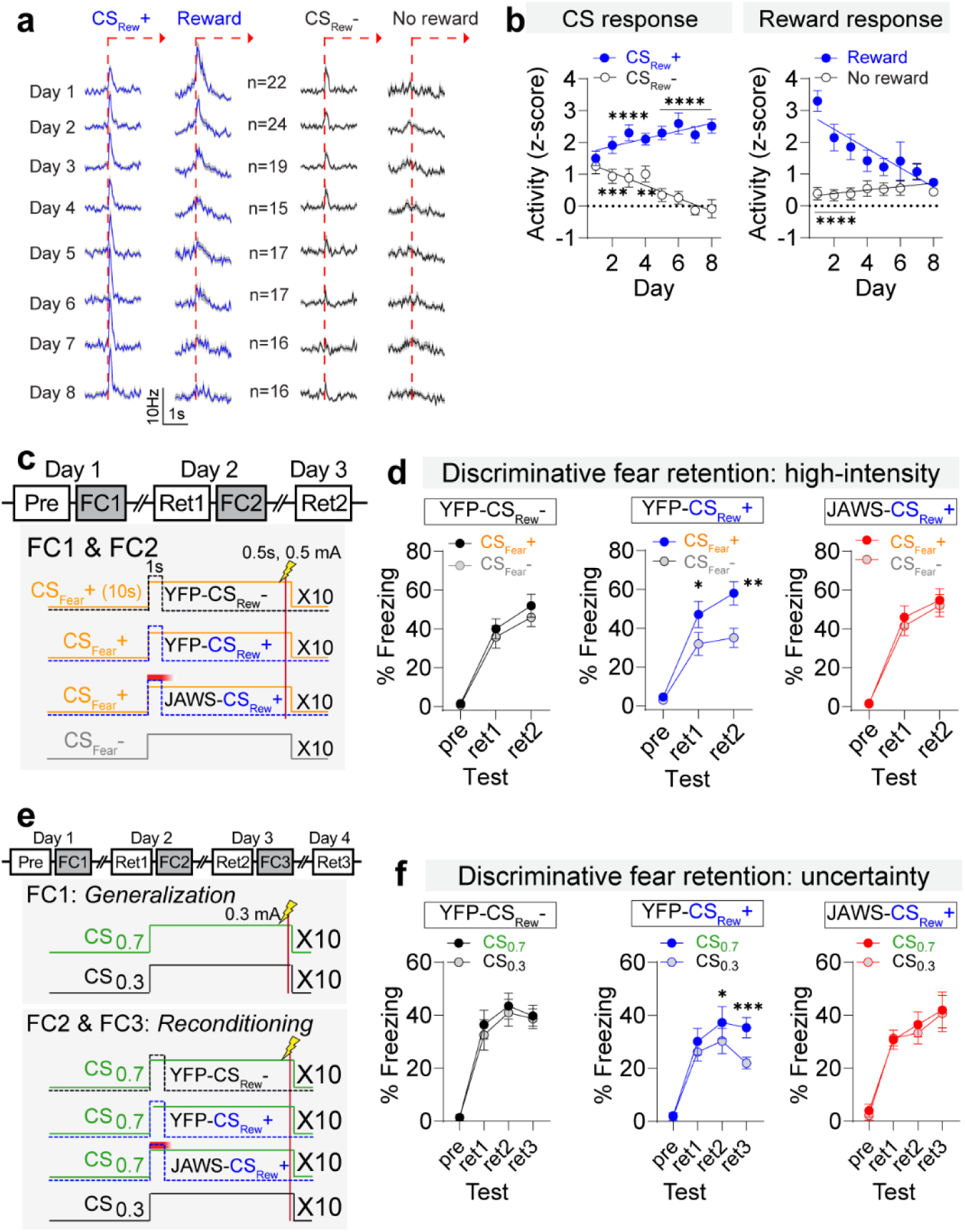
Impact of CS_Rew_+ evoked dopamine on discriminatory fear. (**a**) Average dopaminergic responses to CSs and head entries for reward consumption (total 146 cells from 7 mice). (**b**) Normalized responses to CS response (CS_Rew_+ and CS_Rew_-) and Reward response(reward retrieval and no reward response after CS_Rew_-), relative to baseline firing before each CS. CS_Rew_+ and CS_Rew_- responses were positively and negatively correlated with training days, respectively (Pearson’s correlation, CS_Rew_+, *r* = 0.26, *P* = 0.002 and negatively CS_Rew_-, *r* = -0.43, *P* < 0.001). Reward responses were negatively correlated in CS_Rew_+ trials (*r* = -0.4, *P* < 0.001). Significant differences between CS_Rew_+ versus CS_Rew_-, and Reward versus No reward following the CS_Rew_- were observed (\*\**P* < 0.01, \*\*\**P* < 0.001, \*\*\*\**P* < 0.0001). (**c**) Schematic of timing sequences for CS_Rew_+ and CS_Rew_- presentations and JAWS-mediated inhibition during fear conditioning. (**d**) Freezing responses to the CS_Fear_+ and CS_Fear_- during the pre-test, retention tests 1 and 2. CS_Rew_+ mice showed significant discriminatory freezing (\**P* < 0.05, \*\**P* < 0.01, YFP-CS_Rew_+ = 10 mice, YFP-CS_Rew_- = 10 mice, and JAWS- CS_Rew_+ = 10 mice). (**e**) Schematic of probabilistic fear conditioning paradigm to induce fear generalization (FC1) and CS_Rew_+ reversal (FC2 and FC3). (**f**) Following the induction of a generalized threat response, reconditioning with the CS_Rew_+ facilitated discrimination between the CS_0.7_ and CS_0.3_ that was not observed with co-presentation of the CS_Rew_- or when dopamine neurons were inhibited by JAWS during the CS_Rew_+ presentation (\**P* < 0.05, \*\*\**P* < 0.001, YFP-CS_Rew_+ = 10 mice, YFP- CS_Rew_- = 10 mice, and JAWS- CS_Rew_+ = 10 mice). All data presented as mean ± S.E.M. Detailed information about statistical results is provided in Extended Data Table 1.

Having confirmed the discriminatory encoding of reinforcement, we next assessed whether the presentation of the CS_Rew_+ or CS_Rew_- could influence discriminatory fear learning. To determine whether these effects are dopamine neuron dependent, DAT-Cre mice were injected with a virus for conditional expression of the inhibitory light-activated channel JAWS^23^ (AAV-FLEX-JAWS-EGFP) or control virus (AAV-FLEX-EYFP) and an optical fiber was implanted over the VTA (**Extended Data Fig. 1h,i**). Following positive reinforcement learning, mice underwent discriminatory fear conditioning (**Fig. 1c,d** and **Extended Data Fig. 1j,k**) using a 0.5mA foot shock US, which has been shown to induce generalized threat responses^13^. During fear conditioning, CS_Rew_+ or CS_Rew_- was played for 1 s at the onset of the CS_Fear_+ (10 s tone), but not the non-threat predictive stimulus (CS_Fear_-, 10s tone). In a separate group of mice, VTA dopamine neurons were inhibited during the 1 s of CS_Rew_+ presentation (JAWS-CS_Rew_+) (**Fig. 1c**). Mice presented with the CS_Rew_+ at the onset of the CS_Fear_+ displayed discriminatory fear learning, whereas mice presented with the CS_Rew_-, or had dopamine neurons inhibited during the presentation of the CS_Rew_+ displayed generalized threat responses following conditioning (**Fig. 1d**).

As threat intensity increases, uncertainty also increases as a result of stress-related signaling that promotes generalization^3,13^. To determine whether a positive stimulus can prevent generalization associated with uncertainty, mice were conditioned using a probabilistic conditioning paradigm (**Extended Data Fig. 2a**) that has been shown to induce generalization when threat intensity is relatively low^8,13^. Brief co-presentation of the CS_Rew_+, but not the CS_Rew_-, with the higher probability fear CS (CS_0.7_), promoted discriminatory fear learning that was blocked in mice with inhibition of VTA dopamine neurons during the CS_Rew_+ presentation (**Extended Data Fig. 2b-e**).

We next asked whether an established generalized fear response could be reversed by reconditioning mice with co-presentation of CS_Rew_+. Mice were conditioned using the probabilistic conditioning paradigm to induce generalization, followed by two days of reconditioning with co-presentation of the CS_Rew_+ or CS_Rew_- with the CS_0.7_ (**Fig. 1e** and **Extended Data Fig. 3a-c**). All groups of mice showed generalized fear after a single conditioning session (**Fig. 1f**; ret1), but mice presented with the CS_Rew_+ at the onset of the CS_0.7_ during reconditioning displayed improved discrimination (**Fig. 1f**; YFP-CS_Rew_+; ret2,3). In contrast, mice presented with the CS_Rew_- or had dopamine neurons inhibited by JAWS during the presentation of the CS_Rew_+ displayed persistent generalized threat responses (**Fig. 1F**; YFP**-** CS_Rew_- and JAWS- CS_Rew_+; ret2,3).

### A reward-associated cue can modify fear-associated dopamine release in the CeA

Our results demonstrate that VTA dopamine neurons play an important role in the ability of a stimulus associated with positive reinforcement to modify generalized fear behavior. The CeA has emerged as an important site for regulating discriminatory fear learning^13,24–26^, and dopamine signaling in the CeA is critical for discriminatory fear^27^. However, the dynamics of dopamine release in the CeA during either positive or negative reinforcement remain unknown. To begin to address this, we first assessed whether activation of VTA dopamine neurons evokes dopamine release in the CeA. DAT-Cre mice were injected with a virus to conditionally express the red-shifted stimulatory opsin Chrimson^28^ (AAV-FLEX-Chrimson-tdTomato) in dopamine neurons in the VTA and a virus to express the dopamine sensor dLight1.3b^29^ in the CeA (AAV-dLight1.3b, **Fig. 2a**). Stimulation of dopamine neurons evoked transient dopamine release in a frequency- and stimulus duration-dependent manner (**Fig. 2b,c** and **Extended Data Fig. 3d,e**). Next, we tested whether a CS_Rew_+ modifies dopamine release in the CeA to the CS_0.7_ or CS_0.3_ following the induction of generalization and reconditioning (**Fig. 2d**). Similar to our findings with single-unit recording of VTA dopamine neurons during positive reinforcement learning, dopamine was initially released in the CeA to the US (**Fig. 2e-h**). As learning progressed (**Extended Data Fig. 3f**), dopamine signals emerged to the CS_Rew_+ but not the CS_Rew_- and responses to the reward US diminished (**Fig. 2e-h**). Interestingly, in response to the reward US activation to the reward retrieval (head entry to the food dispenser; HE_Rew_) became smaller and more transient but was followed by the emergence of a robust decrease in dopamine (**Fig. 2g,h**). During probabilistic fear conditioning (**Extended Data Fig. 3g,h**), dopamine release was initially detected in response to the US_Fear_ (**Fig. 2i**; FC1). Co-presentation of the CS_Rew_+ during reconditioning did not alter dopamine release to the fear CS_0.7_ but significantly reduced dopamine release to the CS_0.3_ (**Fig. 2i,j**; FC2 and FC3).

**Figure 2.**
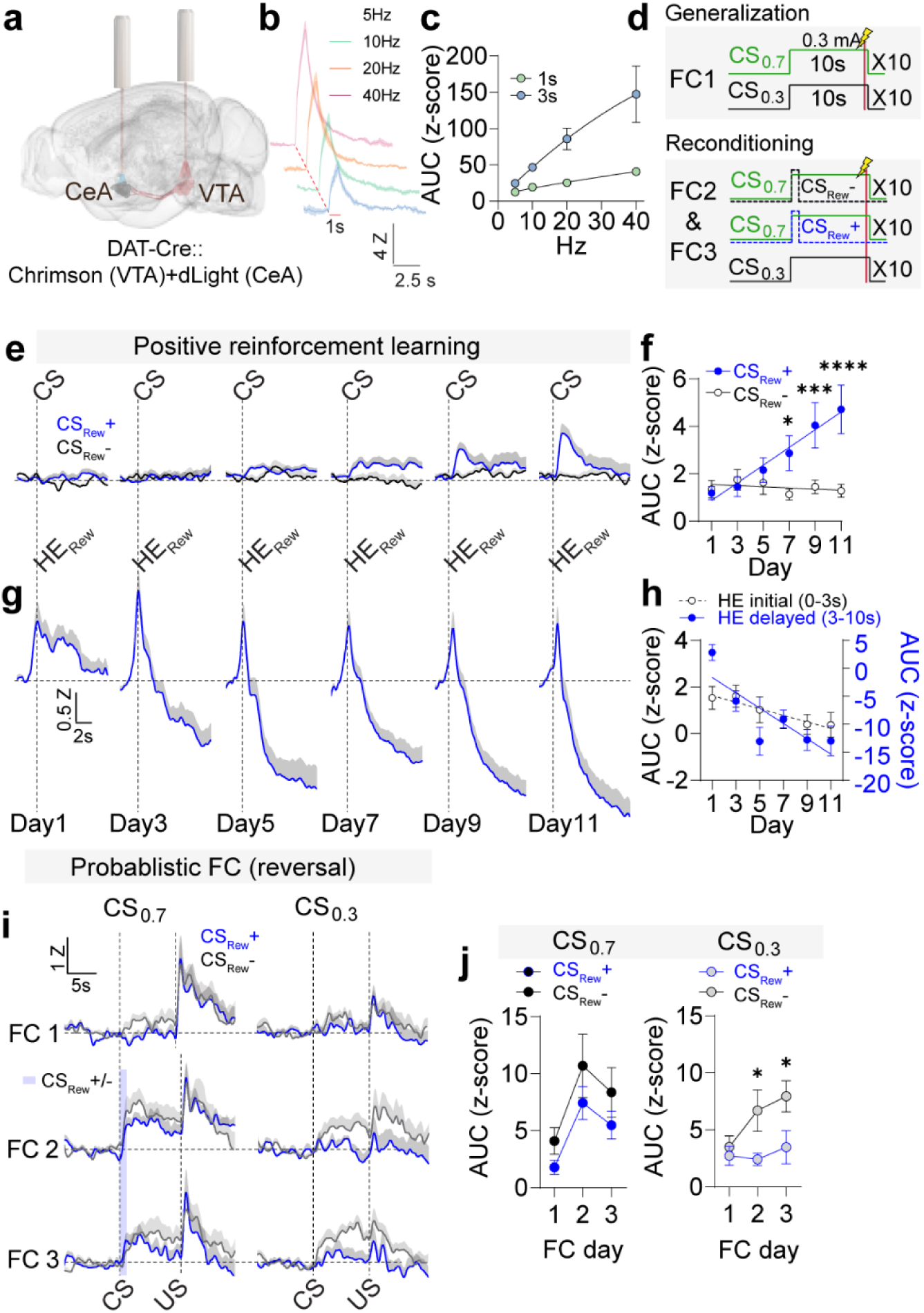
Impact of CS_Rew_+ on fear-evoked dopamine in the CeA. (**a**) Schematic illustrating expression of Chrimson-tdTomato in the VTA and dLight1.3b in the CeA. (**b**) Mean light-evoked dopamine signals in the CeA following VTA stimulation at different frequencies. (**c**) Area under curve (AUC) of dopamine signals to VTA stimulations. (**d**) Schematic of probabilistic fear conditioning paradigm to induce fear generalization and CS_Rew_+ reversal. (**e**) Average traces showing increased dopamine release in the CeA during positive reinforcement learning to the CS_Rew_+ but not to the CS_Rew_- (N = 19 mice). (**f**) Average AUC for CS_Rew_+ and CS_Rew_- (\**P* < 0.05, \*\*\**P* < 0.001, \*\*\*\**P* < 0.001). (**g**) Averaged CeA dopamine signals in response to head entry for reward retrieval (HE_Rew_). (**h**) AUC during head entry for reward retrieval segregated into the initial response (0-3 s, R^2^ = 0.04941, *P* = 0.0147) and secondary response (3-10 s, R^2^ = 0.1953, *P* < 0.0001). (**i**) Dopamine signal in the CeA during fear conditioning with the CS_0.7_ and CS_0.3_ (FC1) and during reconditioning (FC2 and FC3) with the CS_Rew_+ (N = 9) or CS_Rew_- (N = 10). (**j**) Mean AUC of the z-scored dopamine signals in the CeA to the CS_0.7_ and CS_0.3_ during fear conditioning and reconditioning. Co-presentation of the CS_Rew_+ prevented the increase in dopamine release in the CeA in response to the CS_0.3_ (\**P* < 0.05). All data presented as mean ± S.E.M. Detailed information about statistical results is provided in Extended Data Table 1.

### VTA dopamine modifies fear encoding in the CeA

*How does dopamine regulate fear encoding in the CeA*? Subpopulations of genetically distinct CeA neurons are activated (Fear-On) or inhibited (Fear-Off) by conditioned threat stimuli and contribute to associative fear learning^25,26,30–33^. However, how these neurons respond during generalized fear or the impact of dopamine on this encoding is not known. To address this question, we sought to simplify our approach. First, we established whether VTA dopamine projections to the CeA are sufficient to facilitate the reversal of threat generalization. To test this, we expressed ChR2 (AAV-FLEX-ChR2) or a control virus mCherry (AAV-FLEX-mCherry) in VTA dopamine neurons (DAT-Cre mice) and implanted optical fibers over the CeA (**Extended Data Fig. 4a**) to allow for precise stimulation of dopamine terminals for 1 s (20 Hz) during reconditioning (**Extended Data Fig. 4b**). Fear conditioning with a high-intensity US (0.5 mA) induced generalized responses as observed previously. Stimulation of VTA dopamine terminals in the CeA effectively reduced generalized fear responses with just a single reconditioning session (**Extended Data Fig. 4c-e**). Next, we asked whether this stimulation was sufficient to reverse generalization under increased uncertainty, and if so, whether the effect was dopamine-dependent. To achieve this, we expressed ChR2 in VTA dopamine neurons and conditionally mutated the gene encoding tyrosine hydroxylase (*Th*), the rate limiting enzyme in dopamine production, in these same cells using CRISPR/SaCas9 mutagenesis (**Fig. 3a-c**)^34^. Optical fibers were implanted over the CeA (**Fig. 3a** and **Extended Data Fig. 4f**) to deliver 1 s of dopamine terminal stimulations (20 Hz) at the onset of the high probability CS_0.7_ during reconditioning (**Fig. 3d** and **Extended Data Fig. 4g**). Again, stimulation of dopamine terminals effectively induced discrimination between CS_0.7_ and CS_0.3_ in a dopamine-dependent manner (**Fig. 3e,f**).

**Figure 3.**
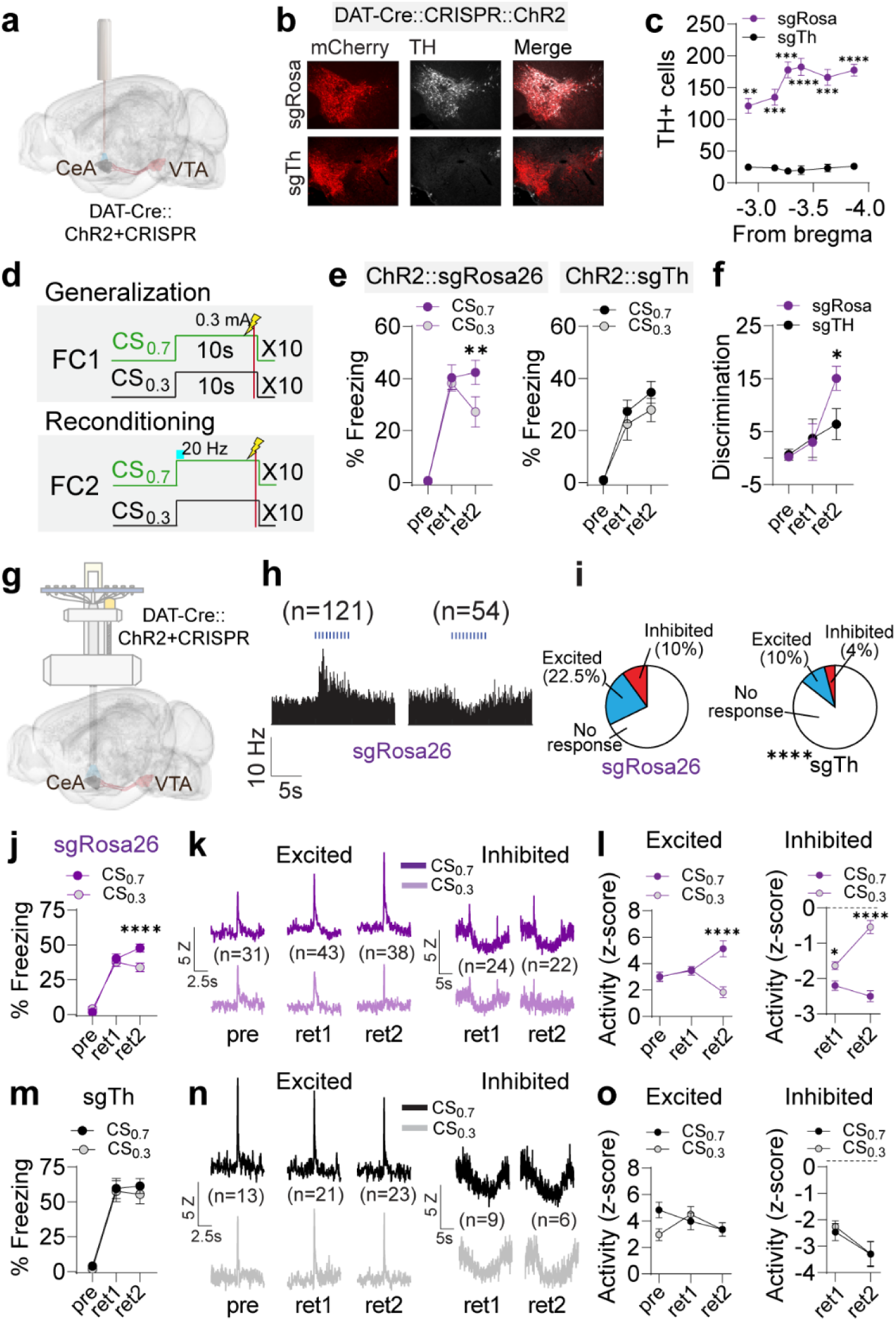
Dopamine release in the CeA is critical for reversing generalization. (**a**) Schematic of AAV-FLEX-ChR2-mCherry and AAV-FLEX-SaCas9-u6-sg*Th* injected into the VTA of DAT-Cre mice and optical fiber placement over the CeA. (**b**) Histological validation of *Th* mutation. (**c**) Quantitative analysis of reduced TH levels in the VTA of sg*Rosa26* control (N = 6) and sg*Th* (N = 5) mice (\*\**P* < 0.01, \*\*\**P* < 0.001, \*\*\*\**P* < 0.001, 6 sections/mice). (**d**) Schematic of probabilistic fear conditioning paradigm and optogenetic stimulation during reconditioning. (**e**) Following the induction of generalized fear response, reconditioning with the optical stimulation of dopamine terminals for 1s facilitated discrimination between the CS_0.7_ and CS_0.3_ that was not observed in *Th* mutagenized mice (\*\**P* < 0.01; sg*Rosa26* = 10 mice and sg*Th* = 10 mice). (**f**) Comparison of fear discrimination between groups (\**P* < 0.05). (**g**) Schematic of AAV-FLEX-ChR2-mCherry and AAV-FLEX-SaCas9-u6-sg*Th* injected into the VTA of DAT- Cre mice and optrode implant for analysis of dopamine neuron activity. (**h**) Examples of excitatory and inhibitory responses to dopamine terminal stimulation from sg*Rosa26* mice. (**i**) Responses of CeA neurons to dopamine terminal stimulation showing reduced excitatory and inhibitory responses in sg*Th* compared to sg*Rosa26* mice. (**j**) Freezing to the CS_0.7_ and CS_0.3_ in sg*Rosa26* mice with CeA recordings during preconditioning, retention test 1, and retention test 2 (\*\*\*\**P* < 0.0001; sg*Rosa26*= 12 mice). (**k**) Average firing to CS_0.7_ and CS_0.3_ in sg*Rosa26* mice during pre-test, retention test 1, and retention test 2. (**l**) Normalized responses to CS_0.7_ and CS_0.3_ (relative to baseline firing before each CS) in sg*Rosa26* mice. During pre-test and retention test 1, there was no discriminatory encoding in excited cells. Following reconditioning with dopamine terminal stimulation, discriminatory encoding was significantly enhanced (\**P* < 0.05, \*\*\*\**P* < 0.0001). In inhibited cells, a small but significant discrimination between the encoding of the CS_0.7_ and CS_0.3_ was observed during retention test 1, but this was greatly enhanced following reconditioning with dopamine terminal stimulation. (**m**) Freezing to the CS_0.7_ and CS_0.3_ in sg*Th* mice with CeA recordings during pre-test, retention test 1, and retention test 2 (sg*Th* = 6 mice). (**n**) Average firing to CS_0.7_ and CS_0.3_ in sg*Th* mice during pre-test, retention test 1, and retention test 2. (**o**) Normalized responses to CS_0.7_ and CS_0.3_ (relative to baseline firing before each CS) in sg*Th* mice. During pre-test, retention test 1 and retention test 2, there was no discriminatory encoding in excited cells. In inhibited cells, no discrimination between the encoding of the CS_0.7_ and CS_0.3_ was observed during retention test 1 or retention test 2. All data presented as mean ± S.E.M. Detailed information about statistical results is provided in Extended Data Table 1.

To assess effects of dopamine signaling on fear encoding in the CeA, DAT-Cre mice were injected with ChR2 and *Th-*CRISPR virus or *Rosa26* control virus in the VTA and implanted an optical fiber and recording electrodes in the CeA (**Fig. 3g** and **Extended Data Fig. 4h**). Stimulation of dopamine-releasing terminals (10 pulses at 20 Hz) in the CeA resulted in both neuronal excitation and inhibition (**Fig. 3h**). The proportion of neurons responsive to dopamine terminal stimulation was significantly reduced in mice with mutated *Th* (**Fig. 3i**). Within our control group (sg*Rosa26*), stimulation of dopamine terminals (1s, 20 HZ) enhanced discriminatory fear following reconditioning (**Fig. 3j** and **Extended Data Fig. 4i,j**), as above. Within the CeA, a subset of neurons was phasically excited by both CSs during the pre-test phase (**Fig. 3k**), but no neurons were detected that were inhibited by either CS (**Fig. 3k** and **Extended Data Fig. 4k**). Following the first conditioning session when mice displayed generalized freezing responses, we observed equivalent transient excitations and prolonged inhibitions (**Fig. 3k,l**; ret1). After reconditioning with dopamine terminal stimulation, we observed discriminatory freezing behavior, which was associated with discriminatory encoding of the CS_0.7_ and CS_0.3_ by CeA neurons in sg*Rosa26* control mice (**Fig. 3k,l**; ret2). In contrast, reconditioning mice with *Th* mutagenesis (sg*Th* group) in VTA dopamine neurons with dopamine terminal stimulation in the CeA did not promote discriminatory behavior (**Fig. 3m**). During the pre-test and first retention test, CeA neurons from sg*Th* mice showed a similar response to those from the sg*Rosa26* group (**Fig. 3n,o** and **Extended Data Fig. 4k**). However, sg*Th* mice had a significantly smaller proportion of inhibited neurons during the retention tests (**Extended Data Fig. 4k**), suggesting that inhibited responses may be mediated in part by dopamine signaling. Following reconditioning, sg*Th* mice exhibited equivalent responses to the CS_0.7_ and CS_0.3_, consistent with their persistent generalized fear responses (**Fig. 3m-o**).

### Differential actions of dopamine receptors in the CeA in the reversal of fear generalization

We now have clear evidence to support a mechanism whereby dopamine can influence discriminatory fear learning, but we do not know which dopamine receptors are responsible for these observed effects. Within the CeA, dopamine D1 and D2 receptors are broadly expressed on subpopulations of inhibitory GABAergic neurons^27,35–37^ where they potently modulate neuronal activity. Related to threat processing, dopamine D2 receptor agonist infused into the CeA can prevent fear generalization, and infusion of dopamine D2 receptor antagonists can promote fear generalization^27^, strongly implicating the importance of this receptor. However, in addition to their ability to signal at the somatodendritic level to modulate neuronal activity, dopamine D2 receptors are also localized to dopamine neuron terminals^38^ and cortical projection neuron terminals^39^; thus, making it difficult to assess the source of D2 receptor that is important for facilitating fear discrimination. To determine whether dopamine receptor expression in the GABA-releasing neurons of the CeA are important for the ability of dopamine terminal stimulation to reverse threat generalization and enhance fear extinction, we generated CRISPR guides to selectively mutate the genes encoding D1 (sg*Drd1*) and D2 (sg*Drd2*) receptors in these neurons. DAT-Cre::VGAT-Flp double transgenic mice were injected with AAV-FLEX-ChR2- mCherry into the VTA and either AAV-FLEXfrt-SaCas9-U6-sg*Drd1*, AAV-FLEXfrt-SaCas9-U6-sg*Drd2* or AAV-FLEXfrt-SaCas9-U6-sg*Rosa26* (control) into the CeA with optical fibers were placed above the bilateral CeA (**Extended Data Fig. 5a,b**). Both sg*Drd1* and sg*Drd2* CRISPRs resulted in significant reductions in mRNA levels in the CeA relative to the sg*Rosa26* control CRISPR (**Fig. 4a,b** and **Extended Data Fig. 5c**). Dopamine receptor mutagenized mice and controls were fear conditioned using the probabilistic conditioning paradigm followed by reconditioning with optical stimulation of dopamine terminals in the CeA (1s, 20Hz) (**Fig. 4c**). Reconditioning facilitated discrimination in the sg*Rosa26* and sg*Drd1* mice, but not in the sg*Drd2* mice, indicating that dopamine signaling mediating the reversal of threat generalization is D2 receptor-dependent (**Fig. 4d** and **Extended Data Fig. 5d,e**). In separate cohorts of mice, we induced fear generalization and reconditioned with co-presentation of the CS_Rew_+ or CS_Rew_- (**Fig. 4e**). The sg*Rosa26* control mice displayed discriminatory fear after the second reconditioning day (**Fig. 4f**). In contrast, the sg*Drd1* group showed discriminatory fear after a single reconditioning, but this effect was not persistent (**Fig. 4f**) and the sg*Drd*2 group did not display discriminatory fear after either reconditioning day.

**Figure 4.**
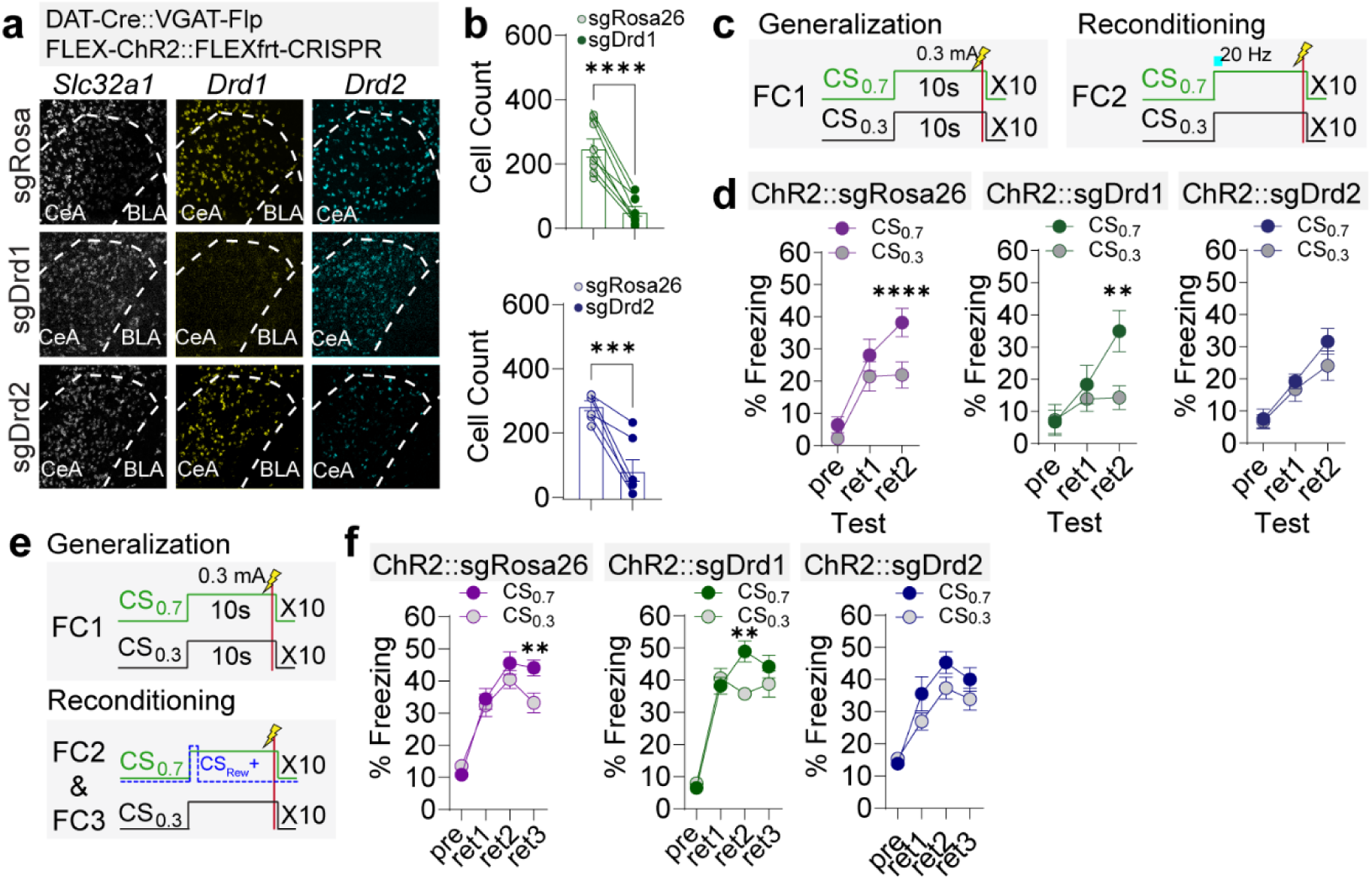
Differential impacts of *Drd1* and *Drd*2 mutagenesis on reversal of generalization. (**a**) RNAscope validation of *Drd1* and *Drd2* mRNA levels in the CeA following mutagenesis. (**b**) Quantitative of reduced mRNA levels associated with nonsense mediated mRNA decay following CRISPR mutagenesis (\*\*\**P* < 0.001, \*\*\*\**P* < 0.0001; sg*Rosa26*/sg*Drd1* = 8 sections from 4 mice and sg*Rosa26*/sg*Drd2* = 7 sections from 4 mice). (**c**) Schematic of probabilistic fear conditioning paradigm and optogenetic stimulation during reconditioning. (**d**) Following the induction of generalized fear response, reconditioning with the optical stimulation of dopamine terminals for 1s facilitated discrimination between the CS_0.7_ and CS_0.3_ in sg*Rosa26* and sg*Drd1* mice that was not observed in *Drd2* mutagenized mice (\*\**P* < 0.01, \*\*\*\**P* < 0.0001, sg*Rosa26* = 15 mice, sg*Drd1* = 10 mice, and sg*Drd2* =16 mice). (**e**) Schematic of probabilistic fear conditioning paradigm to induce fear generalization and CS_Rew_+ reversal. (**f**) Following the induction of generalized fear response, reconditioning with the CS_Rew_+ facilitated discrimination between the CS_0.7_ and CS_0.3_ in sg*Rosa26* and sg*Drd1* mice that was not observed in *Drd2* mutagenized mice (\*\**P* < 0.01, sg*Rosa26* = 16 mice, sg*Drd1* = 9 mice, and sg*Drd2* =14 mice). All data presented as mean ± S.E.M. Detailed information about statistical results is provided in Extended Data Table 1.

Based on the effects of CRISPR mutagenesis of *Drd1* and *Drd2*, we investigated how neurons in the CeA that express these receptors respond to appetitive and aversive stimuli, and how they encode generalized or discriminatory fear. To achieve this, we performed fiber photometry recordings of GCaMP6m *Drd1*-expressing CeA neurons or pro-enkephalin (*Penk*)- expressing neurons (**Fig. 5a** and **Extended Data Fig. 6a**) during appetitive conditioning, fear generalization, and the reversal of generalization (**Fig. 5b**). We chose to utilize *Penk*-Cre mice, as a large proportion of *Drd2-*expressing neurons in the CeA co-express *Penk*^35^ (**Extended Data Fig. 6b,c**), and inactivation of *Penk* impacts fear learning and anxiety^40^. During Pavlovian appetitive conditioning, we observed the emergence of a decrease in calcium in *Drd1* CeA neurons, followed by the emergence of a transient increase at CS_Rew_+ onset (**Fig. 5c,d** and **Extended Data Fig. 6d**). This effect was not observed in *Penk* CeA neurons (**Fig. 5c,d** and **Extended Data Fig. 6d**). In response to head entry and reward retrieval, both populations showed the emergence of an increase in calcium that was slightly larger in the *Penk* CeA population (**Fig. 5e,f**). During fear conditioning, *Drd1* neurons responded to the US but not the CS_0.7_ or CS_0.3_ and reconditioning with the CS_Rew_+ or CS_Rew_- did not affect these responses (**Fig. 5g-i** and **Extended Data Fig. 6e-g**). In contrast, *Penk* CeA neurons responded to the fear US and weakly to the fear CSs during FC1; however, with reconditioning discriminatory responses between the CS_0.7_ or CS_0.3_ emerged in the CS_Rew_+ group but not CS_Rew_- group (**Fig. 5j-l** and **Extended Data Fig. 6e-g**).

**Figure 5.**
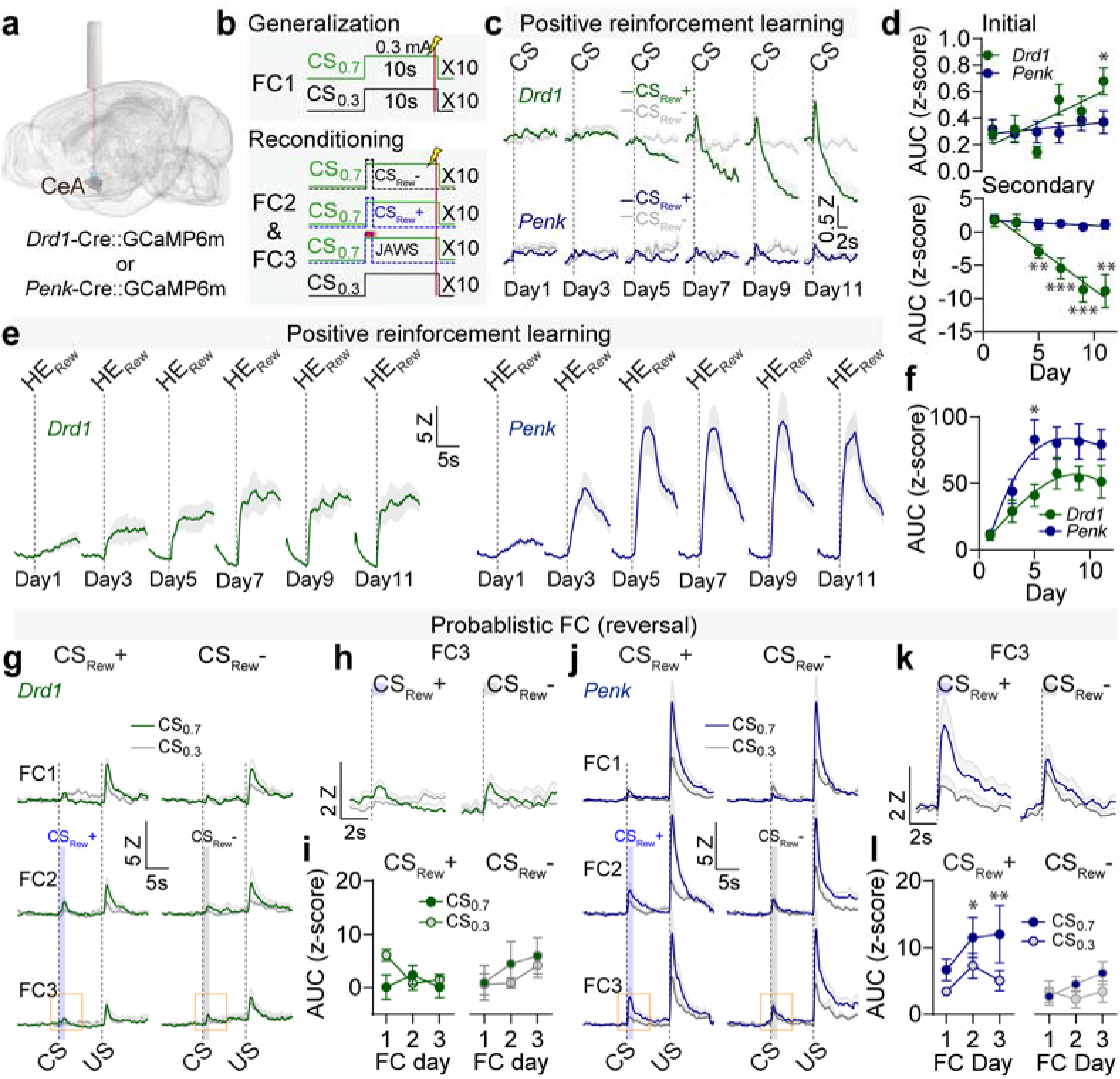
Distinct encoding patterns in *Drd1* and *Penk* CeA neurons during Pavlovian reinforcement learning and reversal of fear generalization. (**a**) Schematic of AAV-FLEX- GCaMP6m injection into the CeA of Drd1-Cre or Penk-Cre mice, with an optical fiber positioned over the CeA. (**b**) Schematic of probabilistic fear conditioning paradigm to induce fear generalization and CS_Rew_+ reversal. (**c**) Average calcium signals from *Drd1* and *Penk* CeA neurons exhibited differential responses to CS_Rew_+ and CS_Rew_-during positive reinforcement learning. (**d**) Area under curve (AUC) for the CS segregated into the initial response (*Drd1*: 0-1 s, R^2^ = 0.1150, *P* = 0.0007) and secondary response (*Drd1*: 2-10 s, R^2^ = 0.3097, *P* < 0.0001). (**e**) Averaged calcium traces of *Drd1* and *Penk* CeA neurons in response to head entry for reward retrieval (HE_Rew_). (**f**) AUC during head entry for reward retrieval (\**P* < 0.05). (**g**) Activity of *Drd1* CeA neurons during fear conditioning with the CS_0.7_ and CS_0.3_ (FC1) and during reconditioning (FC2 and FC3) with the CS_Rew_+ (N = 10 mice) or CS_Rew_- (N = 8 mice). (**h**) Average traces of *Drd1* CeA neurons during the CS presentation in FC3 for CS_Rew_+ and CS_Rew_- groups (magnified view from orange squares in **g**). (i) AUC for the CS across FC1, FC2, and FC3. (**j**) Activity of *Penk* CeA neurons during fear conditioning with the CS_0.7_ and CS_0.3_ (FC1) and during reconditioning (FC2 and FC3) with the CS_Rew_+ (N = 13 mice) or CS_Rew_- (N = 13 mice). (**k**) Average traces of *Penk* CeA neurons during the CS presentation in FC3 for CS_Rew_+ and CS_Rew_- groups (magnified view from orange squares in **j**). (**l**) AUC for the CS across FC1, FC2, and FC3 (* *P* < 0.05, ** *P* < 0.01). All data presented as mean ± S.E.M. Detailed information about statistical results is provided in Extended Data Table 1.

### A reward conditioned stimulus can facilitate fear extinction

During extinction training, dopamine neurons that are inhibited by the CS+ are activated by the omission of the predicted US, generating a type of negative prediction error (NPE) that has been proposed as a type of positive reinforcement^13,41–43^. We found that photosensitive dopamine neurons that are either inhibited or excited by the CS_Fear_+ are activated by omission of the US (foot shock) (**Fig. 6a,b** and **Extended Data Fig. 7a-d**), indicating that these neurons encode both an NPE and the salience of the omitted threat US. To test whether a CS_Rew_+ could also facilitate extinction, mice were conditioned in a simple Pavlovian fear conditioning paradigm and presented with the CS_Rew_+ or CS_Rew_- at the offset of the CS_Fear_+ and CS_Fear_-during extinction training (**Fig. 6c**). Mice presented with the CS_Rew_+ but not the CS_Rew_-, or those with dopamine neurons inhibited during the CS_Rew_+ presentation, displayed enhanced fear extinction and extinction memory recall (**Fig. 6d,e** and **Extended Data Fig. 7e**). The timing of the CS_Rew_+ presentation during the CS_Fear_+ is critical for facilitating extinction, as a random presentation during the intertrial interval did not affect extinction training or extinction memory recall (**Extended Data Fig. 7f-h**).

**Figure 6.**
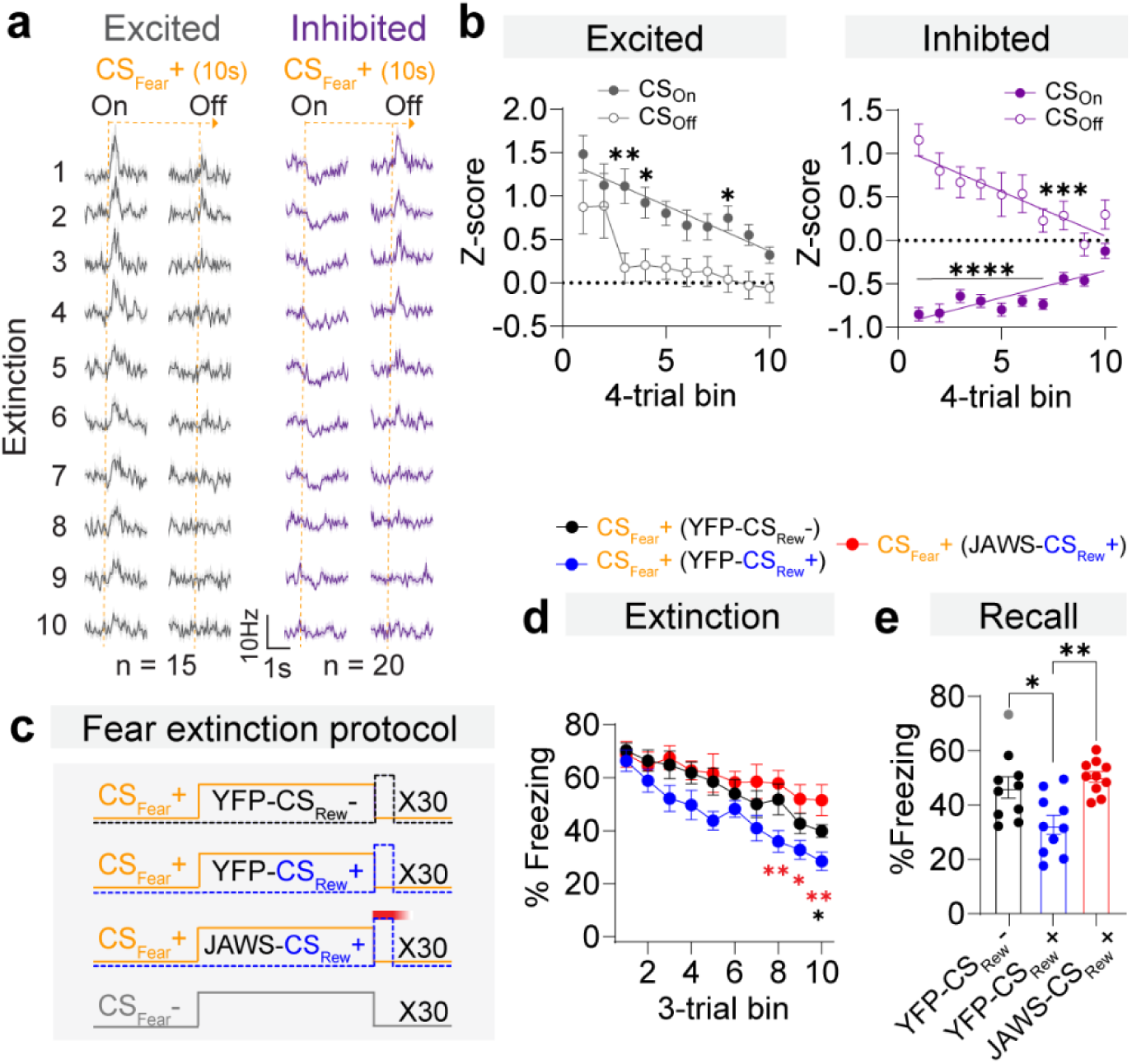
Reward conditioned stimulus enhancement of fear extinction. (**a**) Averaged activity of activated and inhibited neurons extinction trials (3 trial-bin). (**b**) Normalized responses to CS_Fear_+ onset (CS_On_) and CS_Fear_+ offset (CS_Off_) from CS excited neurons (N = 15 cells; CS_On_: R^2^ = 0.1747, *P* < 0.0001, CS_Off_: R^2^ = 0.1021, *P* < 0.0001) and CS inhibited neurons (N = 20 cells; CS_On_: R^2^ = 0.1973, *P* < 0.0001, CS_Off_: R^2^ = 0.1138, *P* < 0.0001). (**c**) Schematic of simple Pavlovian fear conditioning and extinction paradigm with precise timing of CS_Rew_+, CS_Rew_-, and JAWS inhibition during extinction training. (**d**) Fear extinction training in all three groups showing enhanced extinction in the YFP-CS_Rew_+ group (\**P* < 0.05, \*\**P* < 0.01). (**e**) Extinction memory recall showing enhanced recall in the YFP-CS_Rew_+ group (\**P* < 0.05, \*\**P* < 0.01). All data presented as mean ± S.E.M. Detailed information about statistical results is provided in Extended Data Table 1.

Similar to how stimulation of dopamine terminals prevents or reverses fear generalization, brief (1 s) stimulation of dopamine terminals in the CeA at the offset of the CS_Fear_+ and CS_Fear_- (**Fig. 7a**) facilitated extinction and enhanced extinction memory recall in a dopamine-dependent manner (**Fig. 7b,c**). With regard to fear encoding in the CeA, *Th* mutagenized and control mice that had previously undergone fear conditioning and reconditioning (**Fig. 3**) underwent extinction training (**Fig. 7d** and **Extended Data Fig. 7i**). Half of the control mice received dopamine terminal stimulation at the offset of the CS_0.7_ and CS_0.3_, while the other half received no stimulation. All *Th* mutagenized mice received optical stimulation. CeA neurons responsive to the CS were significantly reduced following extinction in mice with dopamine terminal stimulation, compared to non-stimulated controls and *Th* mutagenized mice (**Fig. 7e-g**), indicating that the enhanced extinction observed in the sg*Rosa26* mice that received dopamine terminal stimulation is associated with a greater loss of responsive CeA neurons.

**Figure 7.**
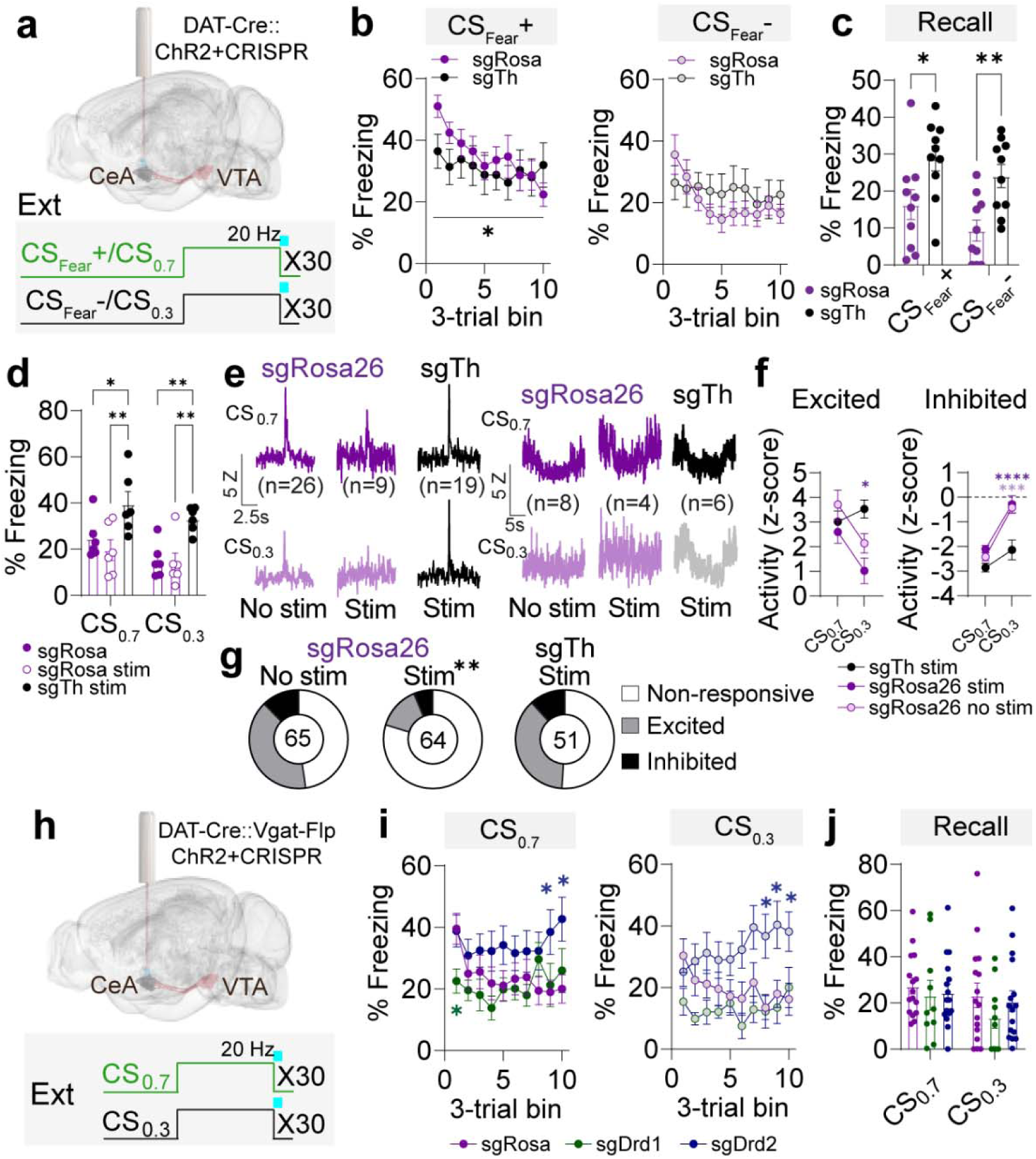
Dopamine dependent extinction of fear encoding in the CeA. (**a**) Schematic of extinction paradigm with precise timing of optical stimulation of dopamine terminals in the CeA during extinction training. **(b**) Fear extinction training in both groups showing enhanced extinction in the sg*Rosa26* stimulated mice (\**P* < 0.05, sg*Rosa26* = 10 mice and sg*Th* = 10 mice). (**c**) Extinction memory recall showing enhanced recall in the sg*Rosa26* stimulated mice (\**P* < 0.05, \*\**P* < 0.01). (**d**) Freezing to the CS_0.7_ and CS_0.3_ during extinction recall with CeA recordings in sg*Rosa26* mice with and without dopamine terminal stimulation and in sg*Th* mice with dopamine terminal stimulation during extinction training (\**P* < 0.05, \*\**P* < 0.01; sg*Rosa26* = 6 mice, sg*Rosa26* stim = 6 mice and sg*Th* stim = 6 mice). (**e**) Average firing to CS_0.7_ and CS_0.3_ in all groups during extinction recall. (**f**) Normalized responses to CS_0.7_ and CS_0.3_ (relative to baseline firing before each CS) in all groups during extinction recall. There were no differences detected in the excited cells in response to the CS_0.7_, but significant differences were observed in response to the CS_0.3_ (\**P* < 0.05, \*\*\**P* < 0.001, \*\*\*\**P* < 0.0001). (**g**) The proportion of cells either excited or inhibited by the CSs during extinction recall was significantly reduced in the sg*Rosa26* mice with dopamine terminal stimulation during extinction training compared to sg*Rosa26* mice without dopamine terminal stimulation and sg*Th* mice with dopamine terminal stimulation (Fisher’s exact test, \*\**P* < 0.01). (**h**) Schematic of extinction paradigm with precise timing of optical stimulation of dopamine terminals in the CeA during extinction training. (**i**) During fear extinction training, freezing to the CS_0.7_ and CS_0.3_ was reduced early in sg*Drd1* mice and increased late in sg*Drd2* mice relative to sg*Rosa26* control mice (\**P* < 0.05). (**j**) Freezing during extinction memory recall was not different between groups. All data presented as mean ± S.E.M. Detailed information about statistical results is provided in Extended Data Table 1.

To assess the roles of *Drd1* and *Drd2* expression, mice underwent extinction training with optical stimulation of dopamine terminals in the CeA (1 s, 20Hz) at the offset of the CS_0.7_ and CS_0.3_ following fear conditioning (**Fig. 7h**). The sg*Rosa26* control mice showed reduced freezing across the extinction session to both CSs (**Fig. 7i**). In contrast, the sg*Drd1* mice displayed reduced freezing from the onset of the extinction training (**Fig. 7i**) and sg*Drd2* mice showed persistent freezing across the extinction session (**Fig. 7i**), indicating that D1 and D2 receptor signaling have opposing actions during extinction training. During extinction recall, all mice showed similar freezing levels (**Fig. 7j**), suggesting that loss of D1 or D2 receptor signaling independently may affect the expression of the extinction responses during training but are not required for the consolidation of the extinction memory.

## Discussion

Our results demonstrate that a stimulus associated with a positive outcome can serve as a potent modulator of fear learning through dopamine-dependent regulation of fear circuitry. We find that the effectiveness of a positive stimulus enhancing fear extinction is dependent on when the CS_Rew_+ is presented, indicating that precise timing is critical. Mechanistically, we demonstrate that generalized fear responses are associated with non-discriminatory encoding in the CeA. Precisely timed transient increases in dopamine within the CeA can facilitate the reversal of generalized fear behavior and non-discriminatory encoding, and the same is true for the facilitation of extinction. The release of dopamine in the CeA in response to both the reward US and fear CS is consistent with this neurotransmitter encoding salience within the structure^14,19,44^.

Within the CeA, *Drd2* expression is most strongly co-localized to *Penk* (>90% of *Drd2^+^* neurons are *Penk* expressing) neurons^27,35–37^. Consistent with our observations that inactivation of *Penk* results in persistent generalized fear following reconditioning with stimulation of dopamine terminals or following reconditioning with CS_Rew_+ presentations, reductions in *Penk* mRNA levels in the CeA are associated with reduced freezing during conditioning and decreased anxiety in rats^40^. In addition to co-expression with *Penk*, *Drd2* is also co-localized with *Prkcd* (PKCδ) expressing neurons^27,35–37^. PKCδ neurons within the CeA have been defined as Fear-Off neurons (inhibited by CS_Fear_+)^32^, and we find significantly fewer CeA inhibited cells following *Th* mutagenesis in the VTA, suggesting that dopamine acting on D2 receptors may be a key mediator of these observed inhibitions. In addition to their responsiveness to fear-associated stimuli^32,45^, PKCδ neurons are also responsive to satiety-related signals^46^ and numerous cells within the CeA have been shown to be responsive to both appetitive and aversive stimuli^44,47,48^. Like the reward consumption-related responses we observed in *Penk-*CeA neurons, *Penk*-expressing neurons of the nucleus accumbens have been shown to be responsive during reward consumption^49^, suggesting a likely broader role for this endogenous opioid in the regulation of food reinforcement. We observed that dopamine release to food reward becomes more inhibited as Pavlovian reward conditioning progresses. In contrast, *Penk*-CeA neurons become progressively more activated, consistent with an inhibitory role for dopamine release on these cells.

*Drd1* is most strongly localized to the medial subdivision of the CeA (CeM), where its expression overlaps with *Tac2*, *Nts*, and *Sst*^35^. The CeM is associated with driving conditioned fear responses^25^, and *Tac2* neurons have been shown to be important for fear memory consolidation^50^, as has dopamine more broadly^51^. Consistent with recent reports that *Sst-*CeA neurons respond to both food and foot shock^44^, we find that *Drd1-*CeA neurons develop conditioned responses to food reward following Pavlovian conditioning and show increasing responses to reward consumption and foot shock; however, these cells do not develop robust conditioned responses to the CS_Fear_+.

We find that inactivation of *Drd1* or *Drd2* in CeA GABA neurons has distinct effects on fear related behaviors. Reduced D1 receptors resulted in the facilitated reduction in freezing during extinction training with VTA dopamine terminal stimulation in the CeA, suggesting that D1 receptor activation promotes the expression of fear memory. In contrast, reduced D2 receptors resulted in persistent freezing during extinction training with VTA dopamine terminal stimulation in the CeA, suggesting that D2 receptor activation may be required for adjusting the certainty of the CS prediction of US delivery in real-time. Inactivation of either receptor did not affect the expression of the extinction memory, suggesting that dopamine signaling in the CeA may not be important for the consolidation of the extinction memory. Instead, dopamine signaling in other regions linked to fear extinction^52^ may be critical for consolidation, such as the medial prefrontal cortex^53,54^, the nucleus accumbens^55,56^ or tail of the striatum^57,58^.

In conclusion, we find that the CeA is a crucial site for integrating information associated with different valences. Stimuli associated with positive reinforcement are an effective means to modify the encoding of CeA neurons and fear-related behavior. Manipulating the salience encoding of dopamine, either naturally or artificially, has a potent influence over discriminatory learning, but the timing is critical.

## References

1 Hovland, C. I. The generalization of conditioned responses. (1937).

2 Dunsmoor, J. E. & Paz, R. Fear Generalization and Anxiety: Behavioral and Neural Mechanisms. Biol Psychiatry 78, 336–343, doi:10.1016/j.biopsych.2015.04.010 (2015).

3 Fellinger, L. et al. A midbrain dynorphin circuit promotes threat generalization. Curr Biol 31, 4388–4396 e4385, doi:10.1016/j.cub.2021.07.047 (2021).

4 Estes, W. K. & Skinner, B. F. Some quantitative properties of anxiety. J Exp Psychol 29, 390–400, doi:DOI 10.1037/h0062283 (1941).

5 Ressler, K. J. et al. Post-traumatic stress disorder: clinical and translational neuroscience from cells to circuits. Nat Rev Neurol 18, 273–288, doi:10.1038/s41582-022-00635-8 (2022).

6 Feng Biao, X. L., Zhang Weixing, Chen Ting, Wang Wenqing, Zheng Xifu. The inhibitive effect of positive emotions on fear generalization. Acta Psychologica Sinica 49, 317–328, doi:10.3724/sp.J.1041.2017.00317 (2017).

7 Zbozinek, T. D. & Craske, M. G. Positive affect predicts less reacquisition of fear: relevance for long-term outcomes of exposure therapy. Cogn Emot 31, 712–725, doi:10.1080/02699931.2016.1142428 (2017).

8 Correia, S. S., McGrath, A. G., Lee, A., Graybiel, A. M. & Goosens, K. A. Amygdala-ventral striatum circuit activation decreases long-term fear. Elife 5, doi:10.7554/eLife.12669 (2016).

9 Janak, P. H. & Tye, K. M. From circuits to behaviour in the amygdala. Nature 517, 284–292, doi:10.1038/nature14188 (2015).

10 Tovote, P., Fadok, J. P. & Luthi, A. Neuronal circuits for fear and anxiety. Nat Rev Neurosci 16, 317–331, doi:10.1038/nrn3945 (2015).

11 Warlow, S. M. & Berridge, K. C. Incentive motivation: ‘wanting’ roles of central amygdala circuitry. Behav Brain Res 411, 113376, doi:10.1016/j.bbr.2021.113376 (2021).

12 Hamati, R., Ahrens, J., Shvetz, C., Holahan, M. R. & Tuominen, L. 65 years of research on dopamine’s role in classical fear conditioning and extinction: A systematic review. Eur J Neurosci 59, 1099–1140, doi:10.1111/ejn.16157 (2024).

13 Jo, Y. S., Heymann, G. & Zweifel, L. S. Dopamine Neurons Reflect the Uncertainty in Fear Generalization. Neuron 100, 916–925 e913, doi:10.1016/j.neuron.2018.09.028 (2018).

14 Steinberg, E. E. et al. Amygdala-Midbrain Connections Modulate Appetitive and Aversive Learning. Neuron 106, 1026–1043 e1029, doi:10.1016/j.neuron.2020.03.016 (2020).

15 Casey, E., Avale, M. E., Kravitz, A. & Rubinstein, M. Dopaminergic innervation at the central nucleus of the amygdala reveals distinct topographically segregated regions. Brain Struct Funct 228, 663–675, doi:10.1007/s00429-023-02614-1 (2023).

16 Watabe-Uchida, M., Zhu, L., Ogawa, S. K., Vamanrao, A. & Uchida, N. Whole-brain mapping of direct inputs to midbrain dopamine neurons. Neuron 74, 858–873, doi:10.1016/j.neuron.2012.03.017 (2012).

17 Schultz, W. Multiple dopamine functions at different time courses. Annu Rev Neurosci 30, 259–288, doi:10.1146/annurev.neuro.28.061604.135722 (2007).

18 Zweifel, L. S. et al. Activation of dopamine neurons is critical for aversive conditioning and prevention of generalized anxiety. Nat Neurosci 14, 620–626, doi:10.1038/nn.2808 (2011).

19 Kong, M. S. & Zweifel, L. S. Central amygdala circuits in valence and salience processing. Behav Brain Res 410, 113355, doi:10.1016/j.bbr.2021.113355 (2021).

20 Backman, C. M. et al. Characterization of a mouse strain expressing Cre recombinase from the 3’ untranslated region of the dopamine transporter locus. Genesis 44, 383–390, doi:10.1002/dvg.20228 (2006).

21 Boyden, E. S., Zhang, F., Bamberg, E., Nagel, G. & Deisseroth, K. Millisecond-timescale, genetically targeted optical control of neural activity. Nat Neurosci 8, 1263–1268, doi:10.1038/nn1525 (2005).

22 Schultz, W., Dayan, P. & Montague, P. R. A neural substrate of prediction and reward. Science 275, 1593–1599, doi:10.1126/science.275.5306.1593 (1997).

23 Chuong, A. S. et al. Noninvasive optical inhibition with a red-shifted microbial rhodopsin. Nat Neurosci 17, 1123–1129, doi:10.1038/nn.3752 (2014).

24 Botta, P. et al. Regulating anxiety with extrasynaptic inhibition. Nature neuroscience 18, 1493–1500, doi:10.1038/nn.4102 (2015).

25 Ciocchi, S. et al. Encoding of conditioned fear in central amygdala inhibitory circuits. Nature 468, 277–282, doi:10.1038/nature09559 (2010).

26 Sanford, C. A. et al. A Central Amygdala CRF Circuit Facilitates Learning about Weak Threats. Neuron 93, 164–178, doi:10.1016/j.neuron.2016.11.034 (2017).

27 De Bundel, D. et al. Dopamine D2 receptors gate generalization of conditioned threat responses through mTORC1 signaling in the extended amygdala. Molecular psychiatry 21, 1545–1553, doi:10.1038/mp.2015.210 (2016).

28 Klapoetke, N. C. et al. Independent optical excitation of distinct neural populations. Nat Methods 11, 338–346, doi:10.1038/nmeth.2836 (2014).

29 Patriarchi, T. et al. Ultrafast neuronal imaging of dopamine dynamics with designed genetically encoded sensors. Science 360, doi:10.1126/science.aat4422 (2018).

30 Duvarci, S., Popa, D. & Pare, D. Central amygdala activity during fear conditioning. The Journal of neuroscience : the official journal of the Society for Neuroscience 31, 289–294, doi:10.1523/JNEUROSCI.4985-10.2011 (2011).

31 Fadok, J. P. et al. A competitive inhibitory circuit for selection of active and passive fear responses. Nature 542, 96–100, doi:10.1038/nature21047 (2017).

32 Haubensak, W. et al. Genetic dissection of an amygdala microcircuit that gates conditioned fear. Nature 468, 270–276, doi:10.1038/nature09553 (2010).

33 Li, H. et al. Experience-dependent modification of a central amygdala fear circuit. Nature neuroscience 16, 332–339, doi:10.1038/nn.3322 (2013).

34 Hunker, A. C. et al. Conditional Single Vector CRISPR/SaCas9 Viruses for Efficient Mutagenesis in the Adult Mouse Nervous System. Cell Rep 30, 4303–4316 e4306, doi:10.1016/j.celrep.2020.02.092 (2020).

35 Kim, J., Zhang, X., Muralidhar, S., LeBlanc, S. A. & Tonegawa, S. Basolateral to Central Amygdala Neural Circuits for Appetitive Behaviors. Neuron 93, 1464–1479 e1465, doi:10.1016/j.neuron.2017.02.034 (2017).

36 McCullough, K. M., Daskalakis, N. P., Gafford, G., Morrison, F. G. & Ressler, K. J. Cell-type-specific interrogation of CeA Drd2 neurons to identify targets for pharmacological modulation of fear extinction. Transl Psychiatry 8, 164, doi:10.1038/s41398-018-0190-y (2018).

37 McCullough, K. M., Morrison, F. G., Hartmann, J., Carlezon, W. A., Jr. & Ressler, K. J. Quantified Coexpression Analysis of Central Amygdala Subpopulations. eNeuro 5, doi:10.1523/ENEURO.0010-18.2018 (2018).

38 Suaud-Chagny, M. F., Ponec, J. & Gonon, F. Presynaptic autoinhibition of the electrically evoked dopamine release studied in the rat olfactory tubercle by in vivo electrochemistry. Neuroscience 45, 641–652, doi:10.1016/0306-4522(91)90277-u (1991).

39 Wang, H. & Pickel, V. M. Preferential cytoplasmic localization of delta-opioid receptors in rat striatal patches: comparison with plasmalemmal mu-opioid receptors. J Neurosci 21, 3242–3250, doi:10.1523/JNEUROSCI.21-09-03242.2001 (2001).

40 Poulin, J. F., Berube, P., Laforest, S. & Drolet, G. Enkephalin knockdown in the central amygdala nucleus reduces unconditioned fear and anxiety. Eur J Neurosci 37, 1357–1367, doi:10.1111/ejn.12134 (2013).

41 Cai, L. X. et al. Distinct signals in medial and lateral VTA dopamine neurons modulate fear extinction at different times. Elife 9, doi:10.7554/eLife.54936 (2020).

42 Salinas-Hernandez, X. I. et al. Dopamine neurons drive fear extinction learning by signaling the omission of expected aversive outcomes. Elife 7, doi:10.7554/eLife.38818 (2018).

43 Felsenberg, J. et al. Integration of Parallel Opposing Memories Underlies Memory Extinction. Cell 175, 709–722 e715, doi:10.1016/j.cell.2018.08.021 (2018).

44 Yang, T. et al. Plastic and stimulus-specific coding of salient events in the central amygdala. Nature 616, 510–519, doi:10.1038/s41586-023-05910-2 (2023).

45 Yu, K. et al. The central amygdala controls learning in the lateral amygdala. Nature neuroscience 20, 1680–1685, doi:10.1038/s41593-017-0009-9 (2017).

46 Cai, H., Haubensak, W., Anthony, T. E. & Anderson, D. J. Central amygdala PKC-delta(+) neurons mediate the influence of multiple anorexigenic signals. Nat Neurosci 17, 1240–1248, doi:10.1038/nn.3767 (2014).

47 Kong, M. S., Ancell, E., Witten, D. M. & Zweifel, L. S. Valence and salience encoding in the central amygdala. Elife 13, doi:10.7554/eLife.101980 (2025).

48 Shabel, S. J. & Janak, P. H. Substantial similarity in amygdala neuronal activity during conditioned appetitive and aversive emotional arousal. Proc Natl Acad Sci U S A 106, 15031–15036, doi:10.1073/pnas.0905580106 (2009).

49 Castro, D. C. et al. An endogenous opioid circuit determines state-dependent reward consumption. Nature 598, 646-+, doi:10.1038/s41586-021-04013-0 (2021).

50 Andero, R., Dias, B. G. & Ressler, K. J. A role for Tac2, NkB, and Nk3 receptor in normal and dysregulated fear memory consolidation. Neuron 83, 444–454, doi:10.1016/j.neuron.2014.05.028 (2014).

51 Fadok, J. P., Dickerson, T. M. & Palmiter, R. D. Dopamine is necessary for cue-dependent fear conditioning. The Journal of neuroscience : the official journal of the Society for Neuroscience 29, 11089–11097, doi:10.1523/JNEUROSCI.1616-09.2009 (2009).

52 Abraham, A. D., Neve, K. A. & Lattal, K. M. Dopamine and extinction: a convergence of theory with fear and reward circuitry. Neurobiol Learn Mem 108, 65–77, doi:10.1016/j.nlm.2013.11.007 (2014).

53 Hikind, N. & Maroun, M. Microinfusion of the D1 receptor antagonist, SCH23390 into the IL but not the BLA impairs consolidation of extinction of auditory fear conditioning. Neurobiol Learn Mem 90, 217–222, doi:10.1016/j.nlm.2008.03.003 (2008).

54 Mueller, D., Bravo-Rivera, C. & Quirk, G. J. Infralimbic D2 receptors are necessary for fear extinction and extinction-related tone responses. Biol Psychiatry 68, 1055–1060, doi:10.1016/j.biopsych.2010.08.014 (2010).

55 Holtzman-Assif, O., Laurent, V. & Westbrook, R. F. Blockade of dopamine activity in the nucleus accumbens impairs learning extinction of conditioned fear. Learn Mem 17, 71–75, doi:10.1101/lm.1668310 (2010).

56 Luo, R. et al. A dopaminergic switch for fear to safety transitions. Nat Commun 9, 2483, doi:10.1038/s41467-018-04784-7 (2018).

57 Menegas, W., Akiti, K., Amo, R., Uchida, N. & Watabe-Uchida, M. Dopamine neurons projecting to the posterior striatum reinforce avoidance of threatening stimuli. Nat Neurosci 21, 1421–1430, doi:10.1038/s41593-018-0222-1 (2018).

58 Chen, A. P. F. et al. Nigrostriatal dopamine modulates the striatal-amygdala pathway in auditory fear conditioning. Nat Commun 14, 7231, doi:10.1038/s41467-023-43066-9 (2023).

