## Supplemental Data for "A neural circuit basis for reward-induced suppression of fear generalization"

**Methods**

**Animals**

Male and female mice aged 2-6 months, including *Slc6a3*^Cre+^ (DAT-Cre), *Slc32a1*^cre+^ (Vgat-Cre), *Drd1a*^Cre+^ (Drd1-Cre), *Penk*^Cre+^(Penk-Cre) and double transgenic mice *Slc6a3*^cre+^::*Slc32a1*^Flp+^ (DAT-Cre::VGAT-Flp) mice were housed on a 12/12 h light/dark cycle (lights on at 7 am) with free access to food and water. Mice were randomly assigned into groups and experiments were conducted during the light cycle. All procedures were performed in compliance with the guidelines set forth by the Institutional Animal Care and Use Committee.

**Virus production and surgery**

All AAV vectors were internally produced (final titer, 1-3 x 10^12^ particles/ml) following previously outlined protocols^1^. Cre-dependent optogenetics viruses included AAV1-FLEX-JAWS-EGFP, AAV1-FLEX-ChR2-mCherry, and AAV1-FLEX-Chrimson-tdTomato and AAV1-FLEX-EYFP (control). The dopamine sensor virus was AAV1-CAG-dLight1.3b. Cre-dependent calcium sensor virus was AAV1-FLEX-GCaMP6m. Cre-dependent CRISPR viruses were AAV1-FLEX-SaCas9-HA-U6-sg*Th,* AAV1-FLEX-SaCas9-HA-U6-sg*Drd1*, AAV1-FLEX-SaCas9-HA-U6-sg*Drd2,* and AAV1- FLEX-SaCas9-HA-U6-sg*Rosa26* (control). Flp-dependent CRISPR viruses were AAV1-FLEXfrt-SaCas9-HA-U6-sg*Drd1*, AAV1-FLEXfrt-SaCas9-HA-U6-sg*Drd2,* and AAV1-FLEXfrt-SaCas9-HA-U6-sg*Rosa26* (control). Viral vectors were stored at -80°C before surgery. On the day of surgery, mice were anesthetized under isoflurane (1-4%) and head fixed in a stereotaxic frame. One of the viral vectors (0.5 µl) was bilaterally or unilaterally injected into the VTA (3.3 mm posterior, 0.5 mm lateral, and 4.5 mm ventral to bregma) or CeA (1.2 mm posterior, 2.9 mm lateral, and 4.6 mm ventral to bregma; for Penk-Cre mice, 0.8 mm posterior was used to target Drd2 co-expressing Penk neurons) at a rate of 0.25 µl/min. Then optic fibers (200 µm inner core, 0.22 NA) and/or a photometry fiber (400 μm fiber, 0.66 NA and 1.25 mm diameter, Doric Lenses) were bilaterally (optic fibers) or unilaterally (fiber photometry fiber; right hemisphere) implanted dorsal to the VTA (4.0 mm ventral to bregma) or CeA (4.2 mm ventral to bregma for optogenetics and 4.7 mm ventral to bregma for fiber photometry). The fibers were secured in place with two anchoring screws and dental cement. For single-unit recording, a microdrive containing 4 tetrodes (25 µm diameter tungsten wire; California Fine Wire) and an optic fiber was implanted dorsal to the VTA (3.7 mm to the brain surface) or CeA (3.8 mm to the brain surface). Tetrodes were cut and gold-plated to reach impedances of 100-300 kΩ tested at 1 kHz. The distance between the fiber and tetrode tips was about 500 µm.

**Pavlovian reinforcement learning**

Mice were calorie-restricted to 85% of their baseline body weight. Each mouse was placed in one of four identical operant boxes (ENV-307W; Med Associates) with a food hopper on one wall, a speaker on the opposite wall, and a smooth plastic panel on the floor. There were 25 rewarding CSs and 25 non-rewarding CSs (CS_Rew_+ and CS_Rew_-; 8 kHz or 15 kHz tone, counterbalanced) trials per day with an average intertrial interval (ITI) of 2 min. The two CSs were presented in a random order. Each CS lasted for 10 s, and only CS_Rew_+ was paired with a 20-mg food pellet (Bio-Serv). As a dependent variable, the number of head entries during the presentation of two CSs was measured using an infrared detector located in the food hopper.

**Fear conditioning and extinction**

Fear conditioning protocols consisted of three tone tests and two conditioning sessions on three consecutive days: pre-conditioning test (pre) in the morning and first fear conditioning in the afternoon on day 1 (FC1), post-conditioning retention test 1 (ret1) and second fear conditioning on day 2 (FC2), and post-conditioning retention test 2 (ret2) on day 3. Standard conditioning sessions were conducted in two identical chambers (ENV-307W; Med Associates), which were located in a different room from where Pavlovian reinforcement learning had been carried out. Each chamber was equipped with two speakers on the opposite walls and 24 shock grids on the floor. A petri dish filled with a 1% acetic acid solution was located under the shock grids (Context B). Animals were placed in the chambers and habituated for 2 min. Then two auditory CSs (CS_Fear_+ and CS_Fear_-; 4 kHz or 12 kHz, counterbalanced) were presented 10 times each in a pseudo-random order with an ITI of 60 s (CS_Fear_+ was always presented on the first trial). CS_Fear_+ (10 s) co-terminated with a 0.5 s footshock (US; 0.3-0.5 mA), but CS_Fear_- (10 s) did not. For the reversal of fear generalization, the four-day paradigm was used: pre and FC1 on day 1 for generalization, ret1 and FC2 on day 2, ret2 and FC3 on day 3, and ret3 on day 4. During FC2 and FC3, either CS_Rew_+ or CS_Rew_- was presented together at the onset of CS_Fear_+ as a compound cue for 1 s to noninvasively activate dopamine neurons. In a separate group, CS_Rew_+ coincided with a continuous red light (640 nm, 5-10 mW, 1-s on followed by 1-s ramp down, LaserGlow) to inhibit JAWS-expressing neurons in the VTA. The chambers were cleaned between animals with the acetic acid solution. To measure fearful responses to both CSs, tone tests were performed in a different environment where white plastic inserts covered the walls and shock grids (context A). Mice were habituated in the inserts for 2 min, and then three CS_Fear_+ and CS_Fear_- trials were presented. For the extinction training, thirty CS_Fear_+ and CS_Fear_- each were delivered in context A. The inserts were wiped with 70% ethanol between animals.

In a probabilistic fear conditioning paradigm, all procedures were the same except for the probability of pairing between CS and US during conditioning sessions. Out of ten sets of alternating CS_0.7_ and CS_0.3_ trials (either 4 kHz or 12 kHz, counterbalanced), seven sets of CS_0.7_ (10 s) were paired with 0.3 mA footshock US in a pseudo-random order. In the other three sets, CS_0.3_ (10 s), but not CS_0.7_, was paired with the footshock US to increase the uncertainty of US prediction (**Extended Data Fig. 2a**). The first set was always assigned as CS_0.7_ -paired and CS_0.3_-unpaired with the US.

To stimulate ChR2-expressing VTA neurons during fear conditioning and extinction training, a pulsed blue light (20 Hz, 473 nm, 1 s, LaserGlow) was delivered at the onset of the CS_0.7_ (probabilistic conditioning) or at the offset of CS_Fear_+ and CS_Fear_- (extinction).

During the tests and conditioning sessions behavior was recorded via a video camera mounted on the ceiling. Movement velocities were calculated using tracking software (Ethovision XT 15, Noldus Technology). Freezing behavior was scored during the presentation of CSs if velocities were less than 0.75 cm/s for at least 1 s. The freezing criterion was determined based on the comparison between automatic and manual scoring using a sample dataset.

**Single-unit recording**

Individual calorie-restricted mice (85% baseline body weight) were placed in a holding cage, and their microdrive was connected to a preamplifier, which transferred neural data to a digital Lynx acquisition system (4SX, Neuralynx). Spiking signals from the tetrodes were amplified, filtered, and digitized at 32 kHz. Unit spikes were recorded for 1 ms when voltage potentials exceeded a predetermined threshold. To identify ChR2-expressing dopamine neurons, 10 blue light pulses (473 nm; 5 ms at 20 Hz) were presented 20 times via the optic fiber of the microdrive. The light intensity was adjusted (5-15 mW/mm^2^) to match light-evoked waveforms and spontaneous ones. Once light-responsive units were found, mice underwent 8 daily recording sessions from the following day. In each session, neuronal basal firing patterns were first measured for 10 min in the holding cage. Then, their firing responses were recorded when mice were engaged in Pavlovian reinforcement learning or fear conditioning test days. After the daily training, mice were transferred back to the holding cage, and 10 light pulses were delivered 20 times. If light-responsive units were not found, all tetrodes were lowered in 80 µm increments until light-responsive units were encountered.

**Fiber photometry**

Calorie-restricted mice were connected to fiber photometry patch cords (Doric Lenses) to record fluorescent dopamine signals (dLight1.3b) or calcium signals from Drd1-expressing neurons and Penk-expressing neurons from the CeA. The dopamine or calcium signals were recorded using RZ5 BioAmp Processor and Synapse software (Tucker Davis Technologies) with a 465 nm LED (531 Hz, sinusoidal, excitation, Doric Lenses) and 405 nm LED (211 Hz, sinusoidal, isosbestic, Doric Lenses). A sampling rate was set at 1,017.25 Hz. The LED intensity was calibrated at the tip of the optic fiber and maintained within the range of 30–40 µW. During behavioral experiments, Med Associates delivered TTL signals associated with CS deliveries and head entries for offline analysis. Dopamine signals were normalized to the baseline (10 s before CSs), and peri-CS activity was computed with a custom script (PEP developed by Dr. Scott Ng-Evans). To validate dopamine release in the CeA by stimulation of the VTA, an optic fiber implanted in the VTA was connected to red light (640 nm, 5 mW, 5 ms pulses, LaserGlow). Stimulation-evoked dopamine signals were recorded in response to diverse light parameters (5, 10, 20, and 40 Hz with 1 s or 3 s duration; 1 min interval, pseudo-random order).

**In situ hybridization**

To validate CRISPR mutagenesis in the CeA, DAT-Cre::VGAT-Flp male and female mice (9 weeks old) were injected with a Flp-dependent sgRNA targeting Drd1 or Drd2 in the left CeA and Rosa26 (control) in the right CeA. Verification was performed using RNAscope (ACDBio RNAscope Multiplex Fluorescent V2). After five weeks, mice were rapidly decapitated, and brains were flash-frozen on dry ice before being stored at −80°C. CeA sections were collected and processed while tracking the right and left hemispheres (25 μm, coronal). Representative sections from injection sites in the CeA were selected for hybridization (AP: −0.83 to −1.67 mm). To quantify Drd2 and Penk mRNA co-expression levels, DAT-Cre mice were similarly decapitated and prepared for sectioning as described above. Representative sections covering the rostral to caudal CeA (AP: −0.6 to −1.8 mm) were collected. RNAscope was performed following the ACDBio V2 protocol. Probes from ACDBio were used to stain for Vgat, Drd1/Drd2, and Cas9 (for CRISPR validation) or Vgat, Drd2, and Penk (for co-expression validation). Slides were coverslipped with Fluoromount containing 4′,6-diamidino-2 phenylindole (DAPI) (Southern Biotech) and imaged using a Leica SP8X confocal fluorescent microscope at the University of Washington Keck Center. Imaging settings were kept consistent across slides for uniform data collection and analysis. Quantification analysis was performed using ImageJ and QuPath 0.4.4 (QuPath).

**Immunohistochemistry**

After the completion of behavioral experiments, mice were euthanized and transcardially perfused with phosphate-buffered saline (PBS) and 4% paraformaldehyde. Brains were extracted and cryoprotected in 30% sucrose in PBS. To confirm virus expression and fiber/electrode placements, frozen brains were sectioned and stained overnight (30 µm for single-unit recording and 50 µm for optogenetics and fiber photometry). The following primary antibodies were used: Rabbit anti-tyrosin hydroxylase (1:1000; Millipore: AB152), Mouse anti-GFP (1:1000; Millipore: MAB3580), Rat anti-mCherry (1:1000; Invitrogen M11217), and Rabbit anti-HA (1:1000; Sigma H6908). Secondary antibodies were used with 1:200 dilution (Jackson ImmunoResearch). Using a Nikon upright microscope and Keyence BZ-X710, images were collected to examine recording sites, fiber placements, and protein expression levels.

**Quantification and statistical analysis**

Single units recorded from the VTA were isolated based on various waveform features using Offline Sorter (Plexon). Only units displaying stable spikes throughout the recording session were further analyzed using Matlab software (MathWorks). To identify dopamine neurons, a cluster analysis was performed based on spike latency (≤ 8 ms) and probability (≥ 0.8) in response to light pulses. The cluster showing high light responsiveness and a high correlation between spontaneous and light-evoked waveforms was considered as dopaminergic. To examine dopamine neuronal activity during Pavlovian reinforcement learning, peri-event time histograms (PETHs; 50 ms bins) were constructed around the time of CSs and rewards. A reward event in each trial was defined as the first head entry into the food hopper after the food delivery. Firing rates in PETHs were transformed to z-scores relative to the pre-CS baseline firing (2.5 s epoch before CS onset). Average dopamine responses to CSs and rewards were measured during the 400 ms window from the onset of each event.

Statistical significance of all data was assessed using Prism software (GrapchPad Prism 10). See Extended Data Table 1 for detailed information about statistical results. Statistical tests for electrophysiological and behavioral results were performed with one-way ANOVAs across groups as well as mixed-design ANOVAs that contained within-subjects factors (e.g., CS and day) and between-subject variables (e.g., group). Once significant interactions were found, suitable post-hoc tests were used to confirm the statistical significance. Pearson’s correlation tests were conducted to establish a relationship between two variables. Two-tailed P values < 0.05 were considered statistically significant. All data were tested for normality and represented as mean ± SEM.

**Acknowledgments:** Supported by National Institutes of Health grants R01DA044315(LSZ), R01MH135538 (LSZ), F32MH127801 (MSK) and National Research Foundation of Korea grant 2022M3E5E8017804 (YSJ). This work was also supported by the University of Washington Center of Excellence in Opioid Addiction Research (P30 DA048736). We thank Dr. James Allen, Dr. Avery Hunker, and Selena Schattauer for assistance with viral production and CRISPR construct design. We thank Dr. Scott Ng-Evans for assistance with photometry analysis. We also thank members of the Zweifel lab for their thoughtful discussion.

**Author contributions:** Y.S.J. and L.S.Z conceptualized the study. M.S.K, Y.S.J., and L.S.Z designed experiments. M.S.K., Y.S.J., G.H.P. and E.S. performed all experiments and collected and analyzed data. M.S.K., Y.S.J. and L.S.Z wrote the paper.

The authors have no competing interests to declare.

**Data availability:** All data associated with this study will be made available upon acceptance of the manuscript.

**Code availability:** Code for fiber photometry analysis was derived from a publicly available source (Tucker Davis Technologies). Code for electrophysiology analysis was generated in house. All code will be available through GitHub (<https://github.com/zweifellab/ephys>).

**References**

1	Gore, B. B., Soden, M. E. & Zweifel, L. S. Manipulating gene expression in projection-specific neuronal populations using combinatorial viral approaches. *Curr Protoc Neurosci* **65**, 4 35 31-20, doi:10.1002/0471142301.ns0435s65 (2013).

**Extended Data**

**Extended Data Table 1.**

| **Figure** | | **Test** | **N** | **Statistics** | **P** | **Posttest** | **p (post)** |
| --- | --- | --- | --- | --- | --- | --- | --- |
| **Main figure statistics** | | | | | | | |
| 1 | b | *CS*  Two-way ANOVA | Day 1=22  Day 2=24  Day 3=19  Day 4=15  Day 5=17  Day 6=17  Day 7=16  Day 8=16 | Interaction: F_(7,138)_=8.496 | P<0.0001 | Sidak’s multiple comparisons | **p<0.01  ***p<0.001  ****p<0.0001 |
|  | b | *Reward*  Two-way ANOVA | Day 1=22  Day 2=24  Day 3=19  Day 4=15  Day 5=17  Day 6=17  Day 7=16  Day 8=16 | Interaction: F_(7,138)_=9.587 | P<0.0001 | Sidak’s multiple comparisons | ****p<0.0001 |
|  | d | Two-way RM ANOVA | N_YFP-CSRew+_ =10 mice | Interaction: F_(22, 18)_ = 37.95 | P<0.0001 | Sidak’s multiple comparisons | *p<0.05  **p<0.01 |
|  | f | Two-way RM ANOVA | N_RewCS+_=10 mice | Interaction: F_(3, 27)_ = 3.369 | P=0.033 | Tukey’s multiple comparisons test | *p<0.05, ***p<0.001 |
| 2 | f | Two-way RM ANOVA | N=19 | Interaction: F_(5, 90)_ = 4.363 | P=0.0013 | Sidak’s multiple comparisons | *p<0.05  **p<0.001  ****p<0.0001 |
|  | j | Two-way RM ANOVA | N_RewCS+_=9 mice; N_RewCS-_=10 mice | Effect of day: F_(2, 34)_ = 3.578  Effect of group: F_(1, 17)_ = 6.372 | Effect of day: P=0.0389, Effect of group: P=0.0218 | Sidak’s multiple comparisons test | *p<0.05 |
| 3 | c | Two-way RM ANOVA | N_sgRosa26_=6 mice; N_sgTh_=5 mice | Interaction: F_(5, 45)_ = 9.190 | P<0.0001 | Sidaks’s multiple comparisons test | **p<0.01, ***p<0.001,  ****p<0.0001 |
|  | e | Two-way RM ANOVA | N_sgRosa26_=10 mice | Interaction: F_(2, 18)_ = 6.834 | P=0.0062 | Tukey’s multiple comparisons | **p<0.01 |
|  | f | Two-way RM ANOVA | N_sgRosa26_=10 mice; N_sgTh_=10 mice | Effect of group: F_(2, 36)_ = 6.886 | P=0.0029 | Sidak’s multiple comparisons test | *p<0.05 |
|  | j | Two-way RM ANOVA | N_sgRosa26_=12 mice | Interaction: F_(2, 22)_ = 20.27 | P<0.0001 | Tukey’s multiple comparisons | ****p<0.0001 |
|  | l | Two-way ANOVA  Excited cells | N_sgRosa26_=12 mice | Interaction:  F_(1, 218)_ = 10.93 | P=0.0011 | Sidak’s multiple comparisons test | ****p<0.0001 |
|  | l | Two-way ANOVA  Inhibited cells | N_sgRosa26_=12 mice | Interaction:  F_­(1, 88)_ = 73.88 | P<0.0001 | Sidak’s multiple comparisons test | *p<0.05, ****p<0.0001 |
| 4 | b | Paired t-test | N_sgDrd1/Rosa26_=4 mice, 8 sections;  N_sgDrd2/Rosa26_=4 mice, 7 sections | sgDrd1/Rosa26 t=9.260, df=7 sgDrd2/Rosa26; t=6.004, df=6 | P<0.0001  P=0.001 |  |  |
|  | d | Two-way RM ANOVA | N_sgRosa26_=15 mice | Interaction:  F_(1.901, 26.61)_ = 4.409 | P=0.0237 | Sidak’s multiple comparisons test | ****p<0.0001 |
|  | d | Two-way RM ANOVA | N_sgDrd1_=10 mice | Interaction:  F_(2, 18)_ = 6.408 | P=0.0079 | Sidak’s multiple comparisons test | ***p<0.001 |
|  | f | Two-way RM ANOVA | N_sgRosa26_=16 mice | Interaction:  F_(2.363, 35.45)_ = 4.193 | P=0.0182 | Sidak’s multiple comparisons test | **p<0.01 |
|  | f | Two-way RM ANOVA | N_sgDrd1_=9 mice | Interaction:  F_(2.193, 17.54)_ = 5.741 | P=0.0105 | Sidak’s multiple comparisons test | **p<0.01 |
| 5 | d | *Initial*  Two-way ANOVA | Data from N_Drd1_=20 and N_Penk_=26 mice; 0-1 s | Interaction:  F_(5, 194)_ = 2.524 | P=0.0307 | Tukey’s multiple comparisons | *p<0.05 |
|  | d | *Secondary*  Two-way ANOVA | Data from N_Drd1_=20 and N_Penk_=26 mice; 2-10 s | Interaction:  F_(5, 194)_ = 12.64 | P<0.0001 | Tukey’s multiple comparisons | **p<0.01, ***p<0.001, |
|  | f |  | N_Drd1_=20 mice N_Penk_=26 mice; | Effect of group:  F_(3.162, 135.3 )_ = 24.11 | P<0.0001 | Tukey’s multiple comparisons | *p<0.05 |
|  | l | *CS_Rew_+* | N_RewCS-_=13 mice; N_RewCS+_=13 mice; | Effect of group:  F_(1,13)_ = 6.646 | P=0.0229 | Tukey’s multiple comparisons | *p<0.05,  **p<0.01 |
| 6 | b | *Activated* | N= 15 cells | Effect of group:  F_(1,14)_ = 52.15 | P<0.0001 | Sidak’s multiple comparisons test | *p<0.05,  **p<0.01 |
|  | b | *Inhibited* | N= 20 cells | Interaction:  F_(9, 171)_ = 8.345 | P<0.0001 | Sidak’s multiple comparisons test | ***p<0.001,  ****p<0.0001, |
|  | d | Two-way RM ANOVA | N_RewCS-_=10 mice; N_RewCS+_=10 mice; N_JAWSCSRew+_= 10 mice | Effect of trial bin: F_(3.971, 107.2)_ = 16.18, Effect of group: F_(2, 27)_ = 6.866 | Effect of trial bin: P<0.0001, Effect of group: P=0.0039 | Tukey’s multiple comparisons test | *p<0.05  **p<0.01 |
|  | e | One-way ANOVA | N_RewCS-_=10 mice; N_RewCS+_=10 mice; N_JAWSCSRew+_= 10 mice | F_(2, 27)_ = 8.148 | P=0.0017 | Dunnett’s multiple comparisons test | *p<0.05, **p<0.01, |
| 7 | b | *CS_Fear_+*  Two-way RM ANOVA | N_sgRosa26_=10 mice; N_sgTh_=10 mice | Interaction:  F_(9, 162)_ = 2.338 | P=0.0167 |  |  |
|  | c | Two-way RM ANOVA | N_sgRosa26_=10 mice; N_sgTh_=10 mice | Effect of group: F_(1, 18)_ = 15.76 | P=0.009 | Sidak’s multiple comparisons test | *p<0.05, **p<0.01 |
|  | d | Two-way RM ANOVA | N_sgRosa26Stim_=6 mice; N_sgRosa26Nostim_=6 mice;  N_sgThStim_=6 mice | Effect of CS:  F_(1, 15)_ = 7.106  Effect of group:  F_(2, 15)_ = 10.43 | Effect of CS:  P=0.0176  Effect of group:  P=0.0014 | Tukey’s multiple comparisons | *p<0.05, **p<0.01 |
|  | f | Two-way ANOVA  Excited cells | N_sgRosa26Stim_=26 cells; N_sgRosa26Nostim_=9 cells;  N_sgThStim_=19 cells | Effect of CS:  F_(1, 38)_ = 6.380 | P=0.0158 | Tukey’s multiple comparisons | **p<0.01 |
|  | f | Two-way ANOVA  Inhibited cells | N_sgRosa26Stim_=8 cells; N_sgRosa26Nostim_=4 cells;  N_sgThStim_=6 cells | Interaction:  F_­(2, 15)_ = 4.625 | P<0.0273 | Tukey’s multiple comparisons | ***p<0.001, ****p<0.0001 |
|  | g | Fisher’s exact test | N_sgRosa26Stim_=64 total cells; N_sgRosa26Nostim_=65  total cells;  N_sgThStim_=51 total cells |  | P = 0.0016 |  |  |
|  | i | *CS_0.7_*  Two-way  RM  ANOVA | N_sgRosa26_=15 mice;  N_sgDrd1_=10 mice; N_sgDrd2_=16 mice | Interaction: F_(18, 351)_ = 1.519 | P=0.0805 | Dunnett’s multiple comparisons test | *p<0.05 |
|  | i | *CS_0.3_*  Two-way  RM  ANOVA | N_sgRosa26_=15 mice;  N_sgDrd1_=10 mice; N_sgDrd2_=16 mice | Interaction:  F_(18, 342)_ = 1.947 | P=0.0121 | Dunnett’s multiple comparisons test | *p<0.05 |
| **Extended data figure statistics** | | | | | | | |
| E1 | g | Independent t-test | N_DA_=146 cells;  N_Non-DA_=121 cells | t=25.88, df=265 | P<0.001 |  |  |
|  | l | Two-way RM ANOVA | N_RewCS+_=10 mice; N_RewCS-_=10 mice; N_JAWSCSRew+_= 10 mice | Effect of day: F_(1.734, 46.82)_ = 7.282, Effect of group: F_(2, 27)_ = 7.583 | Effect of day: P=0.0027, Effect of group: P=0.0024 | Tukey’s multiple comparisons test | *p<0.05, ***p<0.001 |
| E2 | d | Two-way RM ANOVA | N_RewCS+_=10 mice | Interaction: F_(1.943, 17.49)_ = 10.52 | P<0.0001 | Sidak’s multiple comparisons test | **p<0.01 |
|  | e | Two-way RM ANOVA | N_RewCS+_=10 mice; N_RewCS-_=10 mice; N_JAWSCSRew+_= 10 mice | Interaction:  F_(4, 54)_ = 2.514  Effect of day:  F_(1.664, 44.93)_ = 4.458  Effect of group:  F_(2, 27)_ = 4.180 | Interaction:  P = 0.0521  Effect of day:  P = 0.0226  Effect of group:  P = 0.0262 | Tukey’s multiple comparisons test | *p<0.05  **p<0.01 |
| E3 | c | Two-way RM ANOVA | N_RewCS+_=10 mice; N_RewCS-_=10 mice; N_JAWSCSRew+_=10 mice | Effect of group: F_(2, 27)_ = 2.906 | P=0.0791 | Tukey’s multiple comparisons test | *p<0.05 |
|  | g | Two-way RM ANOVA | N_RewCS+_=9 mice; N_RewCS-_=10 mice | Interaction: F _(2.663, 21.30)_ = 10.76 | P=0.002 | Tukey’s multiple comparisons test | *p<0.05 |
|  | h | Two-way ANOVA | N_RewCS+_=9 mice; N_RewCS-_=10 mice | Interaction:  F_(3, 51)_ = 7.124 | P=0.004 | Tukey’s multiple comparisons test | *p<0.05  **p<0.01 |
| E4 | d | Two-way RM ANOVA | N_ChR2_=11 mice; N_mCherryt_=11 mice | Interaction:  F_(2, 20)_ = 8.832 | P = 0.0018 | Tukey’s multiple comparisons test | *p<0.05 |
|  | e | Two-way RM ANOVA | N_ChR2_=11 mice; N_mCherryt_=11 mice | Effect of day: F_(2, 40)_ = 5.392 | P = 0.0084 | Tukey’s multiple comparisons test | *p<0.05 |
|  | k | Fisher’s exact test | N_sgRosa26Stim_=92 Ret1 cells; N_sgThStim_=58 Ret1 cells |  | P = 0.0291 |  |  |
| E5 | c | Paired t-test | N_sgDrd1/Rosa26_ = 4 mice, 8 sections;  N_sgDrd2/Rosa26_ = 4 mice, 7 sections | sgDrd1/Rosa26 t=5.657, df=7 sgDrd2/Rosa26; t=3.828, df=6 | P<0.0008  P=0.0087 |  |  |
|  | e | Two-way RM ANOVA | N_sgRosa26_=15 mice;  N_sgDrd1_=10 mice; N_sgDrd2_=16 mice | Effect of day: F_(2, 76)_ = 12.57 | P<0.0001 | Sidak’s multiple comparisons test | *p<0.05, **p<0.01 |
| E6 | e | Two-way RM ANOVA | N_Drd1_=10 mice | Interaction:  F_(3, 27)_ = 7.056 | P=0.0012 | Sidak’s multiple comparisons test | ****p<0.0001 |
|  | f | Two-way RM ANOVA | N_Penk_=13 mice | Interaction:  F_(3, 36)_ = 3.763 | P=0.0190 | Sidak’s multiple comparisons test | **p<0.01 |
|  | g | *Drd1*  Two-way RM ANOVA | N_RewCS+_=10 mice; N_RewCS-_=8 mice | Effect of day: F_(3, 48)_ = 6.931 | P = 0.0006 | Dunnett’s multiple comparisons test | *p<0.05 |
|  | g | *Penk*  Two-way RM ANOVA | N_RewCS+_=13 mice; N_RewCS-_=13 mice | Interaction:  F_(3, 72)_ = 3.763 | P = 0.0211 | Sidak’s multiple comparisons test | *p<0.05 |
| E7 | c | Independent t-test | N_DA_=124 cells;  N_Non-DA_=100 cells | t=18.14, df=222 | P<0.001 |  |  |

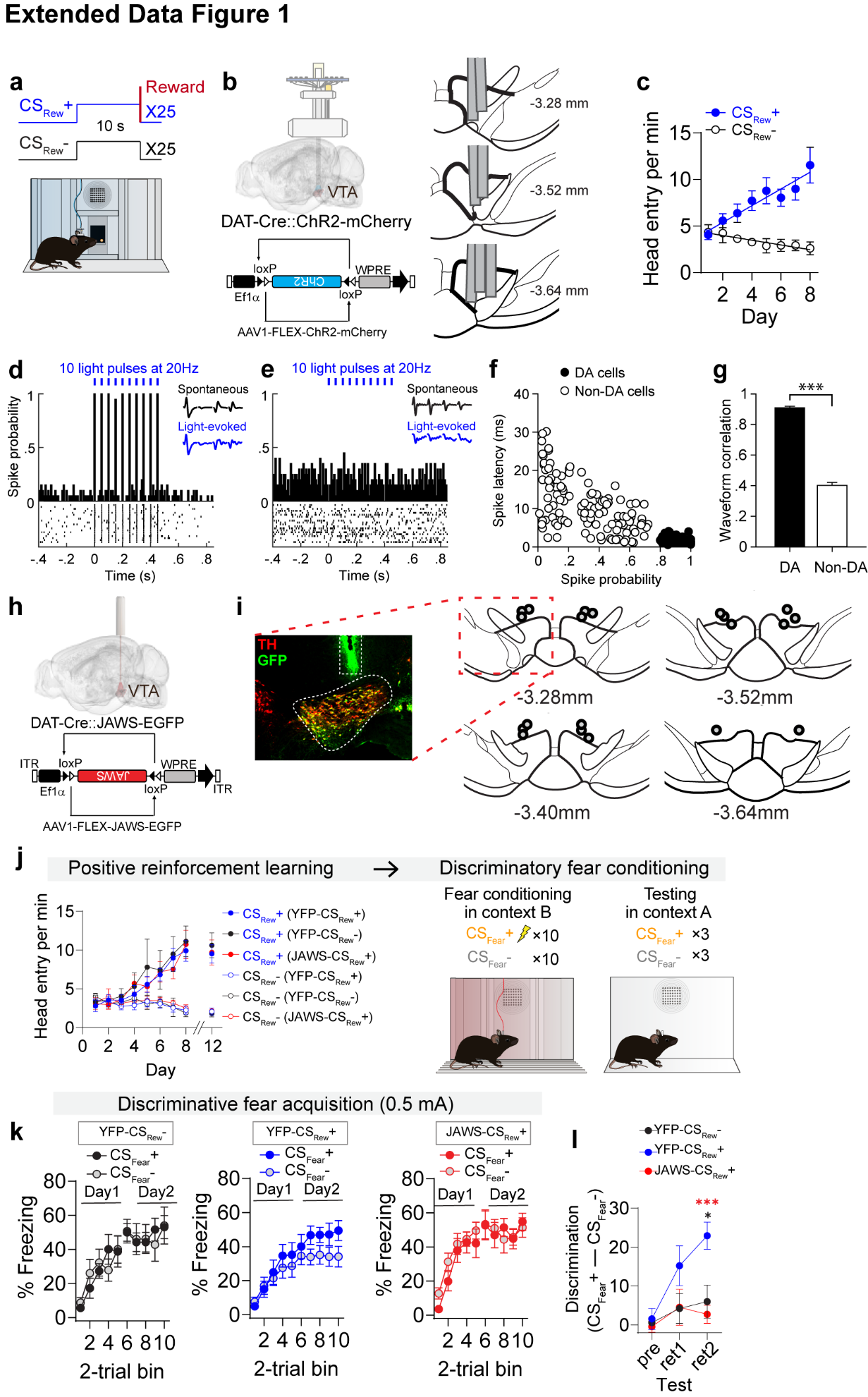

**Extended Data Figure 1. Characteristics of ChR2-responsive dopamine neurons and behavioral results of fear generalization and discrimination.** (**a**) Schematic of positive reinforcement conditioning paradigm. (**b**) Left: Schematic of AAV-FLEX-ChR2-mCherry injected into the VTA of DAT-Cre mice and optrode recording for analysis of dopamine neuron activity. Right: Reconstruction of tetrode tracks included in data analysis. (**c**) Discriminatory cue-evoked head entry behavior in response to CS_Rew_+ and CS_Rew_-. (**d**) Example of VTA neurons displaying light-evoked responses (10 pulses at 20 Hz; 473 nm). (**e**) Example of VTA neurons not showing light-evoked responses. (**f**) Cluster analysis with spike probability and spike latency in response to light. Dopamine neurons are indicated by black circles (spike probability ≥ 0.8 and spike latency ≤ 8 ms). (**g**) Correlations between spontaneous and light-evoked waveforms of dopamine neurons (DA) and non-dopamine neurons (non-DA) (****P* < 0.001). (**h**) Schematic of AAV-FLEX-JAWS-EGFP injected into the VTA of DAT-Cre mice and fiber placement. (**i**) Optic fiber placements in the VTA, along with a representative co-expression of JAWS-GFP and Tyrosine hydroxylase (TH). (**j**) Pavlovian reward conditioning responses in all three groups of mice showing discriminatory cue-evoked head entry responses. Mice were reassessed the day after the final fear conditioning test session to ensure discriminatory reward responding persisted (day 12). Following Pavlovian reward conditioning, mice underwent discriminatory fear conditioning. (**k**) Probabilistic fear acquisition in all three groups (YFP-CS_Rew_- = 10 mice, YFP-CS_Rew_+ = 10 mice, and JAWS-CS_Rew_+ = 10 mice). (**l**) Comparison of fear discrimination between groups (YFP-CS_Rew_+ vs. YFP-CS_Rew_-, **P* < 0.05; YFP-CS_Rew_+ vs. JAWS-CS_Rew_+, ****P* < 0.001). All data presented as mean ± S.E.M. Detailed information about statistical results is provided in Extended Data Table 1.

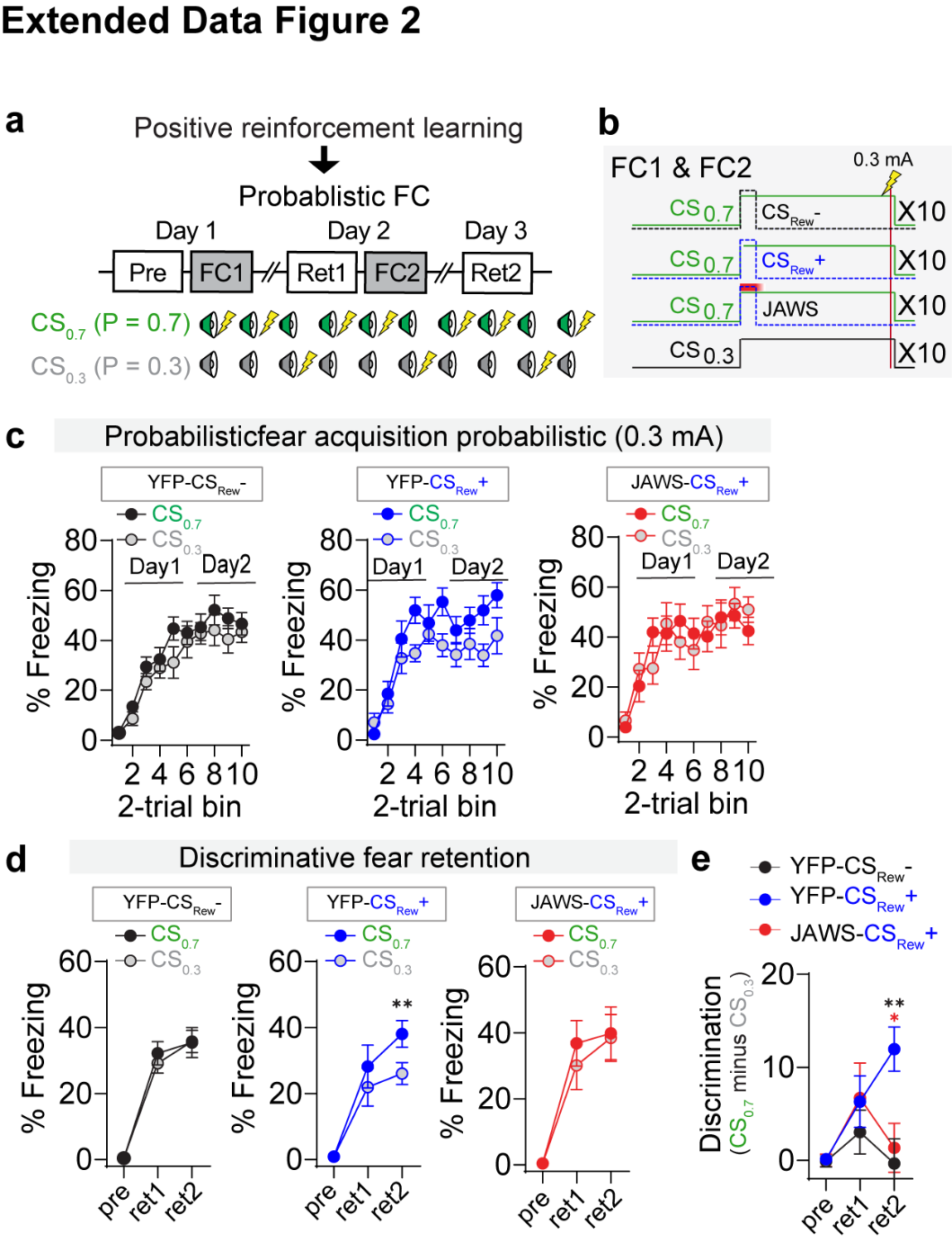

**Extended Data Figure 2. Effects of VTA dopamine neuron inhibition during probabilistic Pavlovian fear conditioning.** (**a**) Schematic of probabilistic Pavlovian fear conditioning. (**b**) Schematic of probabilistic Pavlovian fear conditioning with precise timing of CS_Rew_+, CS_Rew_-, and JAWS inhibition during fear conditioning. (**c**) Discriminative fear acquisition in all three groups with CS_Fear_+ and CS_Fear_- (YFP-CS_Rew_- = 10 mice, YFP-CS_Rew_+ = 10 mice, and JAWS-CS_Rew_+ = 10 mice). (**d**) Discriminative fear retention between CS_0.7_ and CS_0.3_ in all three groups (***P* < 0.01). (**e**) Comparison of fear discrimination between groups (YFP-CS_Rew_+ vs. YFP-CS_Rew_-, ***P* < 0.01; YFP-CS_Rew_+ vs. JAWS-CS_Rew_+, **P* < 0.05). All data presented as mean ± S.E.M. Detailed information about statistical results is provided in Extended Data Table 1.

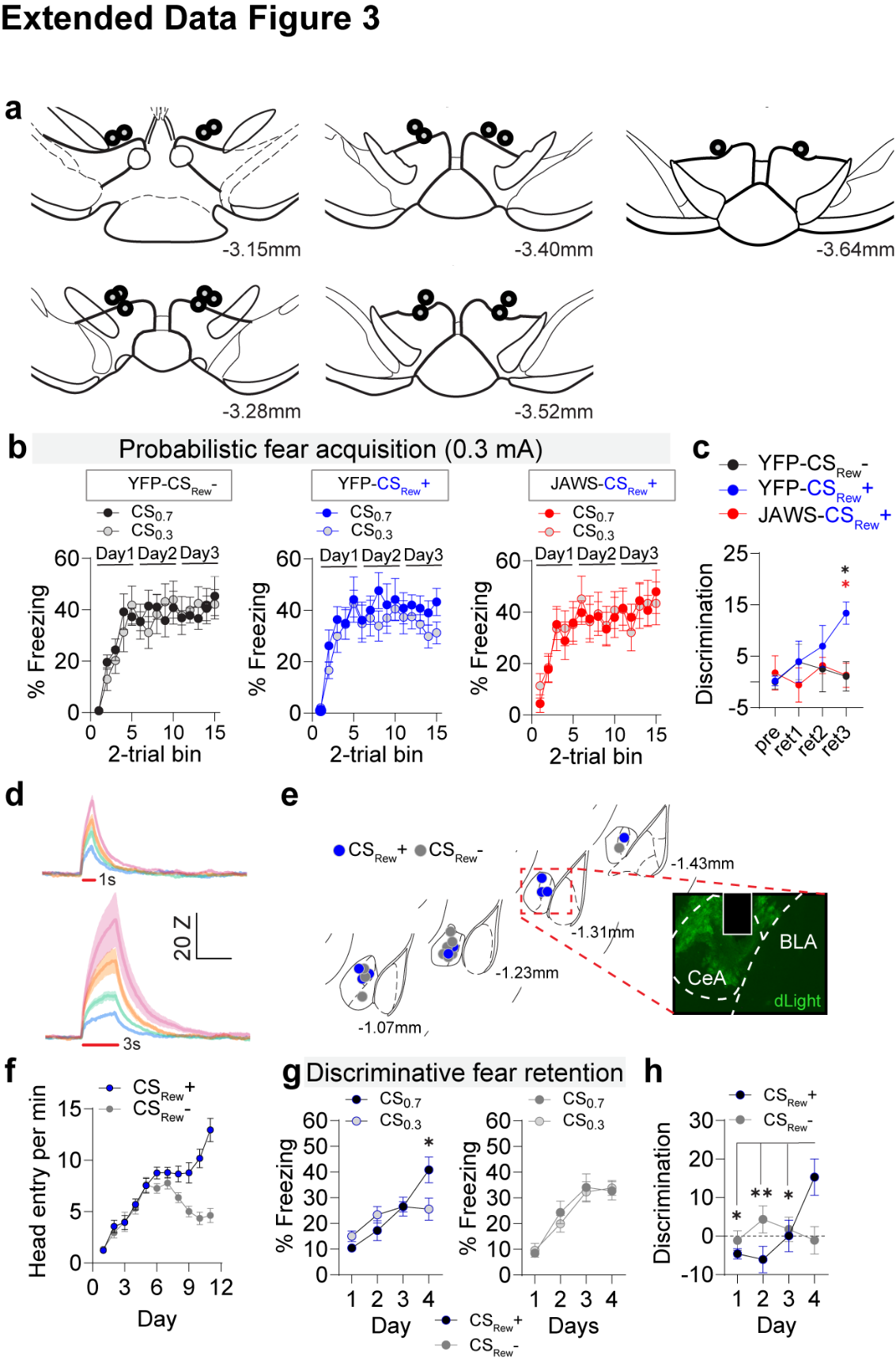

**Extended Data Figure 3. Discriminative fear conditioning and dopamine signals in the CeA.** (**a**) Optic fiber placements in the VTA. (**b**) Acquisition of probabilistic fear conditioning (YFP-CS_Rew_+ = 10 mice, YFP-CS_Rew_- = 10 mice, and JAWS- CS_Rew_+ = 10 mice). (**c**) Discriminative fear retention between CS_0.7_ and CS_0.3_ in all three groups (**P* < 0.05). (**d**) Averaged dopamine signals in the CeA in response to 1 s (top) and 3 s (bottom) of 5 Hz, 10 Hz, 20 Hz, and 40 Hz VTA cell body stimulations (4 mice; 5 times per each stimulation parameter). (**e**) Photometry fiber placements in the CeA (CS_Rew_+ = 9 mice and CS_Rew_- = 10 mice) and a representative expression of dLight. (**f**) The number of head entries per minute during CS_Rew_+ and CS_Rew_- (N = 19 mice). (**g**) Following the induction of a generalized threat response, co-presentation of CS_Rew_+ at the onset of CS_0.7_ induced discriminative fear retention but no effects with CS_Rew_- co-presentation (**P* < 0.05; CS_Rew_+ = 9 mice and CS_Rew_- = 10 mice). (**h**) Comparison of fear discrimination between groups (**P* < 0.05, ***P* < 0.01). All data presented as mean ± S.E.M. BLA, basolateral amygdala. Detailed information about statistical results is provided in Extended Data Table 1.

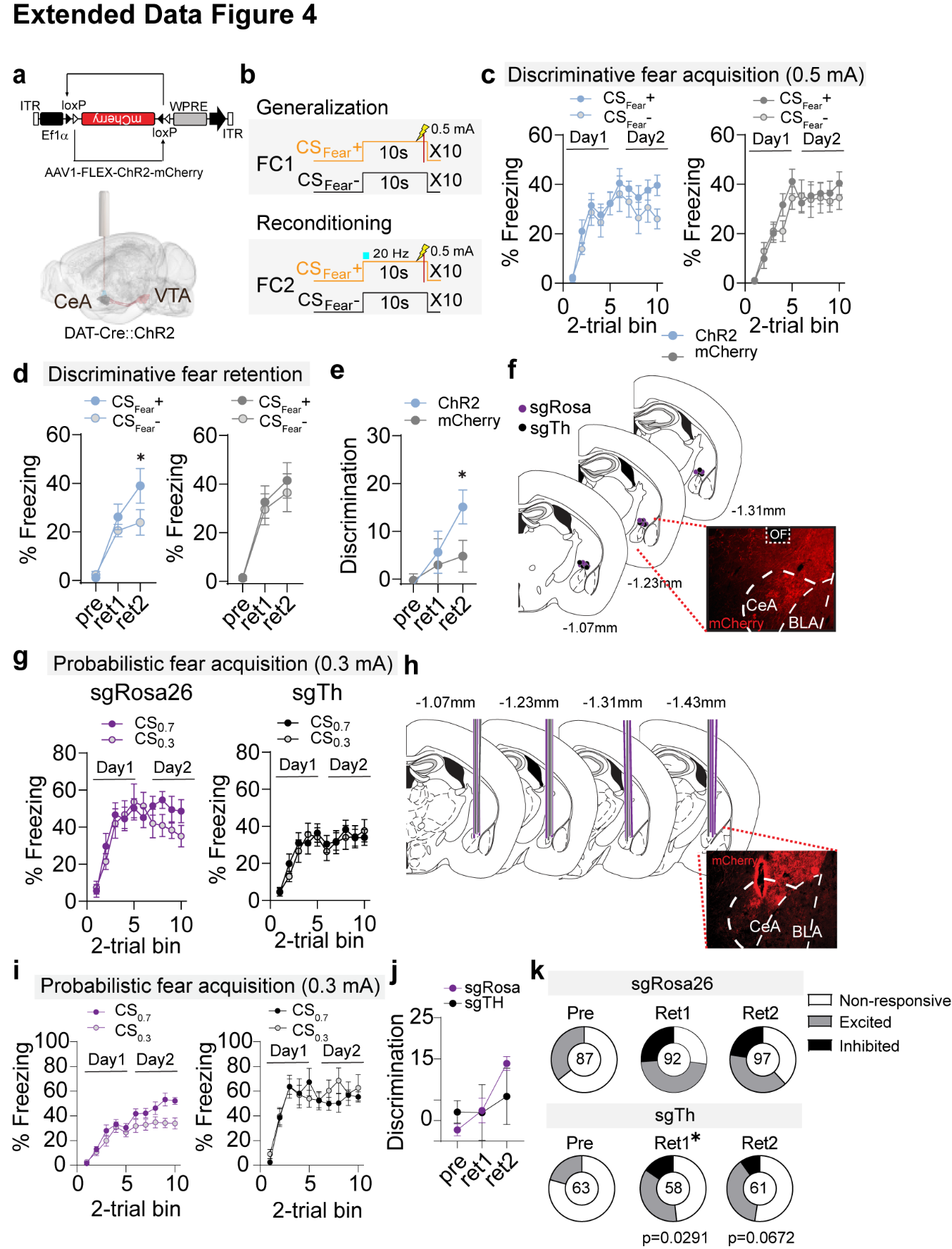

**Extended Data Figure 4. Role of the dopamine input from the VTA to the CeA in discriminative fear conditioning.** (**a**) Schematic of AAV-FLEX-ChR2-mCherry injected into the VTA of DAT-Cre mice and optical fiber placement over the CeA. (**b**) Schematic of discriminative fear conditioning paradigm and optogenetic stimulation during reconditioning. (**c**) Acquisition of discriminative fear conditioning (ChR2 = 11 mice and mCherry = 11 mice). (**d**) Following the induction of a generalized threat response, reconditioning with the optical stimulation of dopamine terminals in the CeA facilitated discrimination (**P* < 0.05). (**e**) Comparison of fear discrimination (**P* < 0.05). (**f**) Optic fiber placements in the CeA and representative ChR2-mChery terminal expression in the CeA. (**g**) Acquisition of probabilistic fear conditioning (sg*Rosa26* = 12 mice and sg*Th* = 6 mice). (**h**) Schematic of AAV-FLEX-ChR2-mCherry and AAV-FLEX-SaCas9-u6-sg*Th* injected into the VTA of DAT-Cre mice and optrode implant for analysis of dopamine neuron activity. Representative VTA terminal expression in the CeA and optrode placements. (**i**) Acquisition of probabilistic fear conditioning. (**j**) Comparison of fear discrimination between groups (group effect: **P* < 0.05).

(**k**) Proportion of non-responsive, excited, and inhibited neurons during pre-test, retention 1, and retention 2 between sg*Rosa26* and sg*Th*. Numbers inside of pie charts indicate the total number recorded (**P* < 0.05). All data presented as mean ± S.E.M. Detailed information about statistical results is provided in Extended Data Table 1.

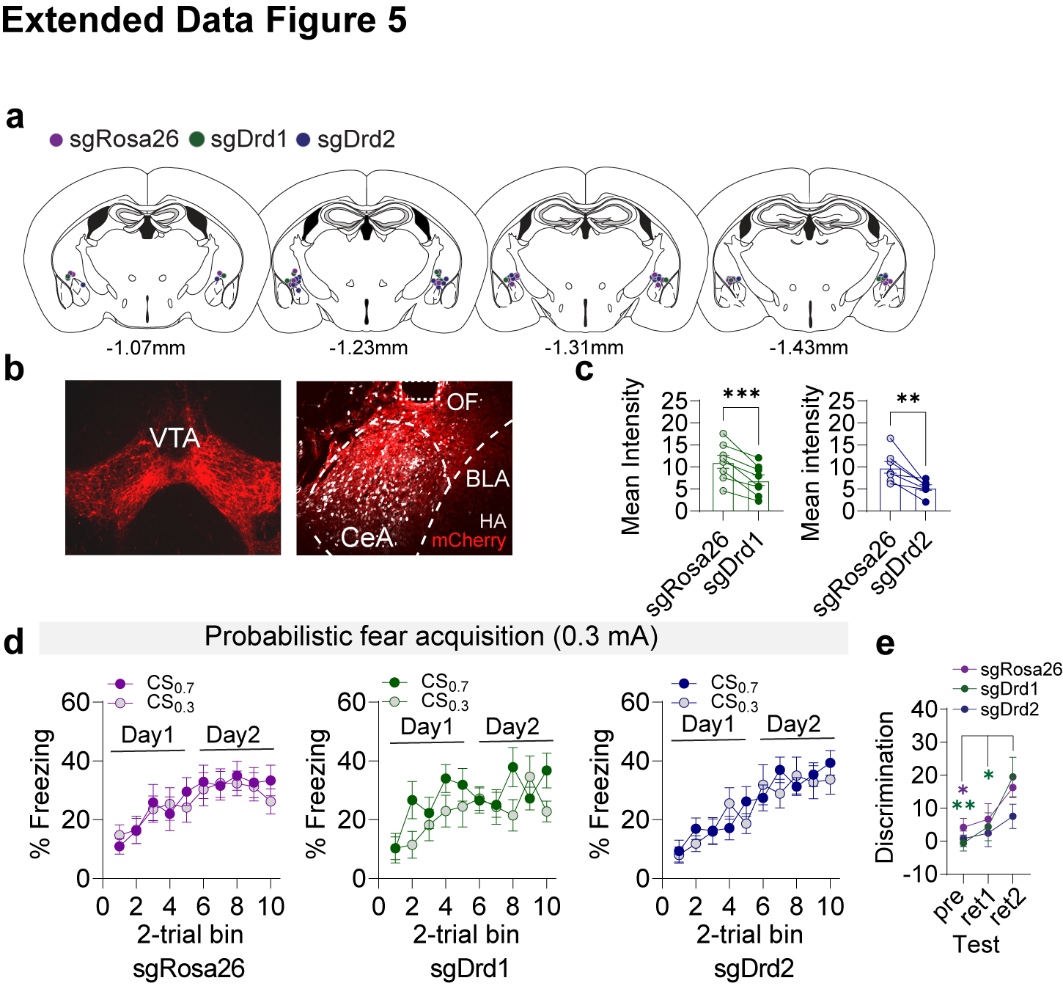

**Extended Data Figure 5. Mutagenesis of *Drd1* and *Drd*2 in the CeA and the acquisition of probabilistic fear conditioning.** (**a**) Optic fiber placements in the bilateral CeA. (**b**) Representative expression of ChR2-mCherry in the VTA (left) and ChR2 terminals in the CeA with HA staining showing CRISPR virus expression (right). (**c**) Comparison of the mean intensity between the control side (sg*Rosa26*) vs. knockout side (sg*Drd1*, left; sg*Drd2,* right; ***P* < 0.01, ****P* < 0.001; sg*Rosa26/Drd1* = 8 sections from 4 mice and sg*Rosa26/Drd2* = 7 sections from 4 mice). (**d**) Acquisition of probabilistic fear conditioning with VTA terminal stimulation in the CeA across all three groups. (**e**) Comparison of fear discrimination between groups (**P* < 0.05, ***P* < 0.01). All data presented as mean ± S.E.M. BLA, basolateral amygdala. OF, optic fiber. Detailed information about statistical results is provided in Extended Data Table 1.

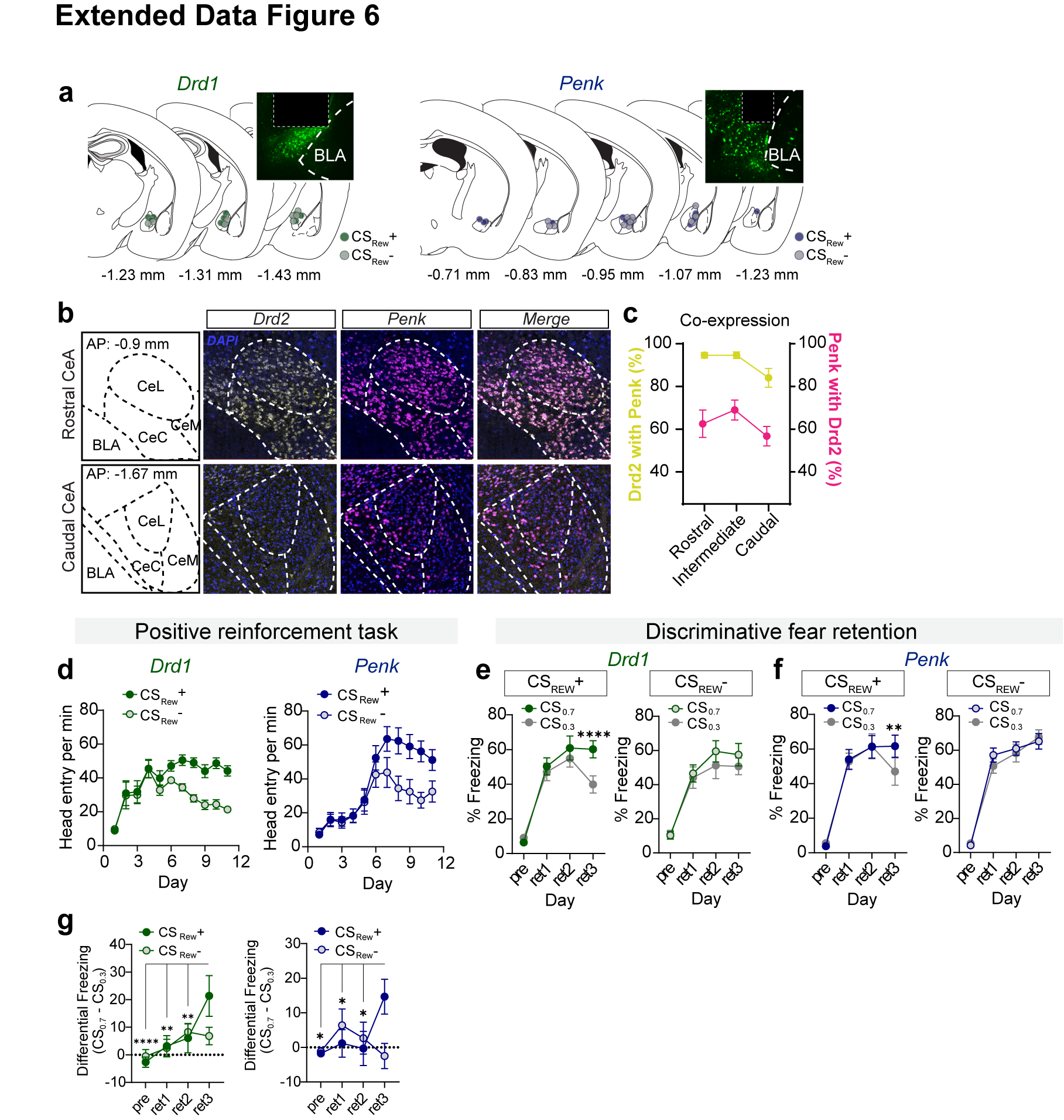

**Extended Data Figure 6. Pavlovian reinforcement learning and reversal of fear generalization in Drd1-Cre and Penk-Cre mice.** (**a**) Optic fiber placements in the CeA of Drd1-Cre (left) and Penk-Cre mice (right). (**b**) RNAscope validation of *Drd2* and *Penk* mRNA co-expression levels in the rostral CeA (top) and caudal CeA (bottom). (**c**) *Drd2* CeA neurons with *Penk* co-expression (yellow) and *Penk* CeA neurons with Drd2 co-expression (pink) across the rostral, intermediate, and caudal CeA. (**d**) The number of head entries per minute during CS_Rew_+ and CS_Rew_- in Drd1-Cre (N = 18) and Penk-Cre (N = 26) mice. (**e**) Following the induction of a generalized threat response, co-presentation of CS_Rew_+ at the onset of CS_0.7_ induced discriminative fear retention but no effects with CS_Rew_- co-presentation in Drd1-Cre mice (*****P* < 0.0001; CS_Rew_+ = 10 mice and CS_Rew_- = 8 mice). (**f**) Following the induction of a generalized threat response, co-presentation of CS_Rew_+ at the onset of CS_0.7_ induced discriminative fear retention but no effects with CS_Rew_- co-presentation in Penk-Cre mice (***P* < 0.01; CS_Rew_+ = 13 mice and CS_Rew_- = 13 mice). (**g**) Comparison of fear discrimination between groups (**P* < 0.05, ***P* < 0.01, *****P* < 0.0001). All data presented as mean ± S.E.M. BLA, basolateral amygdala. Detailed information about statistical results is provided in Extended Data Table 1.

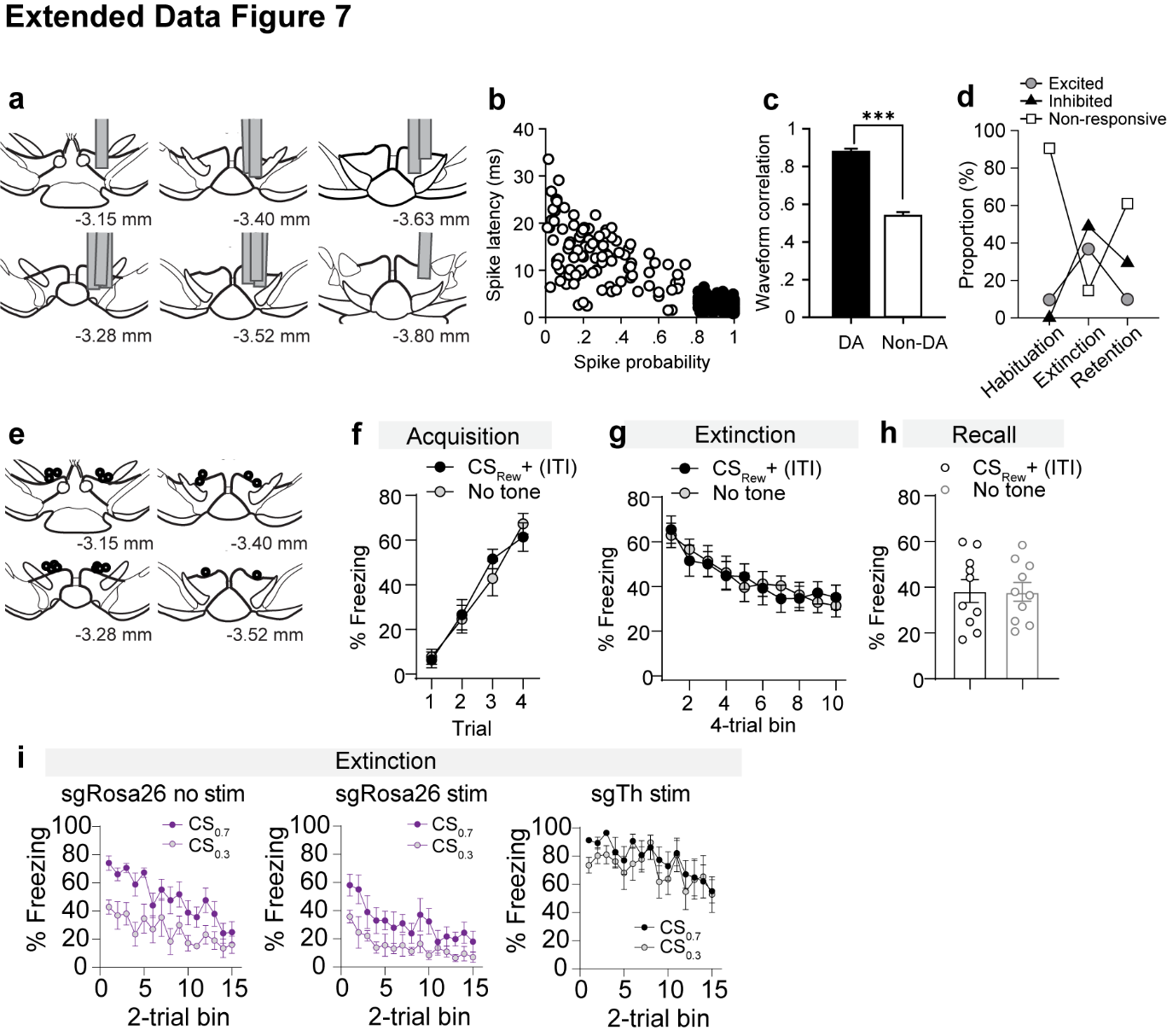

**Extended Data Figure 7. VTA dopamine neuron responses during extinction.** (**a**) Reconstruction of tetrode tracks included in data analysis. (**b**) Cluster analysis with spike probability and spike latency in response to light. Dopamine neurons are denoted by black circles (spike probability ≥ 0.8 and spike latency ≤ 8 ms). (**c**) Correlations between spontaneous and light-evoked waveforms of dopamine neurons (DA) and non-dopamine neurons (non-DA) (****P* < 0.001). (**d**) Proportion of excited, inhibited, and non-responsive neurons during habituation, extinction, and retention. (**e**) Optic fiber placements in the VTA. (**f**) Fear acquisition with mice receiving random CS_Rew_+ presentations (CS_Rew_+ ITI) or no CS_Rew_+ (No tone) presentation (CS_Rew_+ ITI = 10 mice and No tone = 10 mice). (**g**) Extinction training of CS_Rew_+ (ITI) and No tone groups. (**h**) Extinction retention of CS_Rew_+ (ITI) and No tone groups. (**i**) Extinction training of all three groups analysis (sg*Rosa26* no stim = 6 mice, sg*Rosa26* stim = 6 mice, and sg*Th* = 6 mice). All data presented as mean ± S.E.M. Detailed information about statistical results is provided in Extended Data Table 1.
